# ATF6 aggravates angiogenesis-osteogenesis coupling during ankylosing spondylitis by mediating FGF2 expression in chondrocytes

**DOI:** 10.1101/2020.10.08.332379

**Authors:** Mengjun Ma, Hongyu Li, Peng Wang, Wen Yang, Rujia Mi, Yuhang Jiang, Yixuan Lu, Xin Shen, Pengfei Sui, Yanfeng Wu, Huiyong Shen

## Abstract

Although angiogenesis-osteogenesis coupling is important in ankylosing spondylitis (AS), therapeutic agents targeting the vasculature remain elusive. Here, we identified activating transcription factor 6 (ATF6) as an important regulator of angiogenesis in AS patients. Firstly, we found that ATF6 and fibroblast growth factor 2 (FGF2) levels were higher in SKG mice and AS patient cartilage. The pro-angiogenic ability of human chondrocytes was enhanced through activated ATF6-FGF2 axis following long-term stimulation with inflammatory factors, e.g. TNF-α, IFN-γ or IL-17.

Mechanistically, ATF6 interacted with the *FGF2* promotor and promoted its transcription. Treatment with the ATF6 inhibitor Ceapin-A7 inhibited angiogenesis *in vitro* and angiogenesis-osteogenesis coupling *in vivo*. ATF6 may aggravate angiogenesis-osteogenesis coupling during AS by mediating FGF2 transcription in chondrocytes, implying that ATF6 represents a promising therapeutic target for AS.

## Introduction

Ankylosing spondylitis (AS) is a rheumatoid disease characterized by chronic and repeated inflammation, primarily affecting the axial skeleton and joints.^1^ Nearly 40% of patients progress to functional deformities, including joint deformities and rigid spines.^2^ Previous studies have shown that anti-TNF-α treatment is beneficial for AS patients in relieving signs and symptoms and improving physical functions, but it has no effect on delaying radiographic progression.^3–5^ Some researchers hypothesized that the process of pathologic bone formation was independent of inflammation as AS progressed.^6^ However, recent studies showed that long-term treatment with anti-TNF-α for 8 years delayed radiographic progression in AS patients.^7,8^ A similar result was observed in a clinical study on anti-IL-17 treatment in AS patients.^9^ These results indicate that inflammation does play an important role in pathologic bone formation in AS patients.^10^ Further study on the relation between the process of pathological bone formation and inflammation is needed.

Cartilage degradation and replacement by granulation tissue containing vessels is considered an important pathological change before joint ankylosis during AS.^1,11^ The normal cartilage of an adult is avascular due to the presence of many factors that inhibit angiogenesis, including antiangiogenic protease inhibitors and chondromodulin-I (ChM-I), in the extracellular matrix.^12^ However, previous studies showed that the amount of collagen was lower in the articular cartilage of AS patients than in non-AS individuals, indicating that the barrier function of cartilage to vessels might be altered in AS patients.^13^ This kind of change might allow endothelial cells, osteoblasts and other cells to invade into cartilage, followed by granulation tissue formation and active osteogenesis.^14,15^ Thus, to interfere with pathologic osteogenesis in AS, it is necessary to determine how other cell types, such as endothelial cells, invade cartilage.

Numerous studies have shown that endoplasmic reticulum stress (ERS) has a significant regulatory effect on angiogenesis.^16^ ERS is a self-protective response that mainly stabilizes the function of the endoplasmic reticulum by inducing the unfolded protein response (UPR).^17,18^ Among the 3 UPR pathways, the roles of the inositol-requiring enzyme 1 a (IRE1a) and protein kinase RNA-like endoplasmic reticulum kinase (PERK) pathways in angiogenesis are clear.^16^ However, the role of the activating transcription factor 6 (ATF6) pathway in angiogenesis has not been fully elucidated. Ghosh et al.^19^ interfered with ATF6 in the hepatocellular carcinoma cell line HepG2, observed a significant decrease in vascular endothelial growth factor (VEGF) expression, and proved that ATF6 could bind to the promoter region of VEGF. ATF6 is important in cartilage development,^20,21^ but whether ATF6 regulates cartilage by regulating angiogenesis is unclear.

In this study, we show that ATF6 aggravates angiogenesis-osteogenesis coupling during AS by mediating FGF2 expression in chondrocytes. Our results indicate that ATF6 represents a promising therapeutic target for AS.

## Results

### Chondrocytes in AS-associated chronic inflammation increase the expression of FGF2

Whether chondrocytes in patients with AS undergo functional changes after long-term inflammation remains unclear. Previous studies showed that proinflammatory factors such as TNF-α, IFN-γ and IL-17 were significantly increased in the serum and joint fluid of patients with AS.^22^ Thus, we extracted human chondrocytes from hip joints and identified them by IF staining of collagen II (Supplementary Fig. 1). We then stimulated chondrocytes with TNF-α, IFN-γ and IL-17 to observe the changes in angiogenic factor expression (Fig. 1a). After TNF-α stimulation of chondrocytes for 1 day (short-term stimulation) or 7 days (long-term stimulation), the expression of various angiogenic factors was upregulated (Fig. 1b&c). However, short-term stimulated chondrocytes tested 7 days after stimulation showed that the expression of angiogenic factors decreased significantly, close to the level before TNF-α stimulation (Fig. 1b). Moreover, long-term stimulated chondrocytes still expressed high levels of FGF2 and VEGF (Fig. 1c). Similarly, chondrocytes stimulated long-term with IFN-γ or IL-17 also expressed high levels of FGF2 that lasted for 7 days (Fig. 1d). WB and ELISAs showed that FGF2 expression was elevated in chondrocytes after long-term stimulation with TNF-α, IFN-γ or IL-17 (Fig. 1e&f).

**Figure 1.**
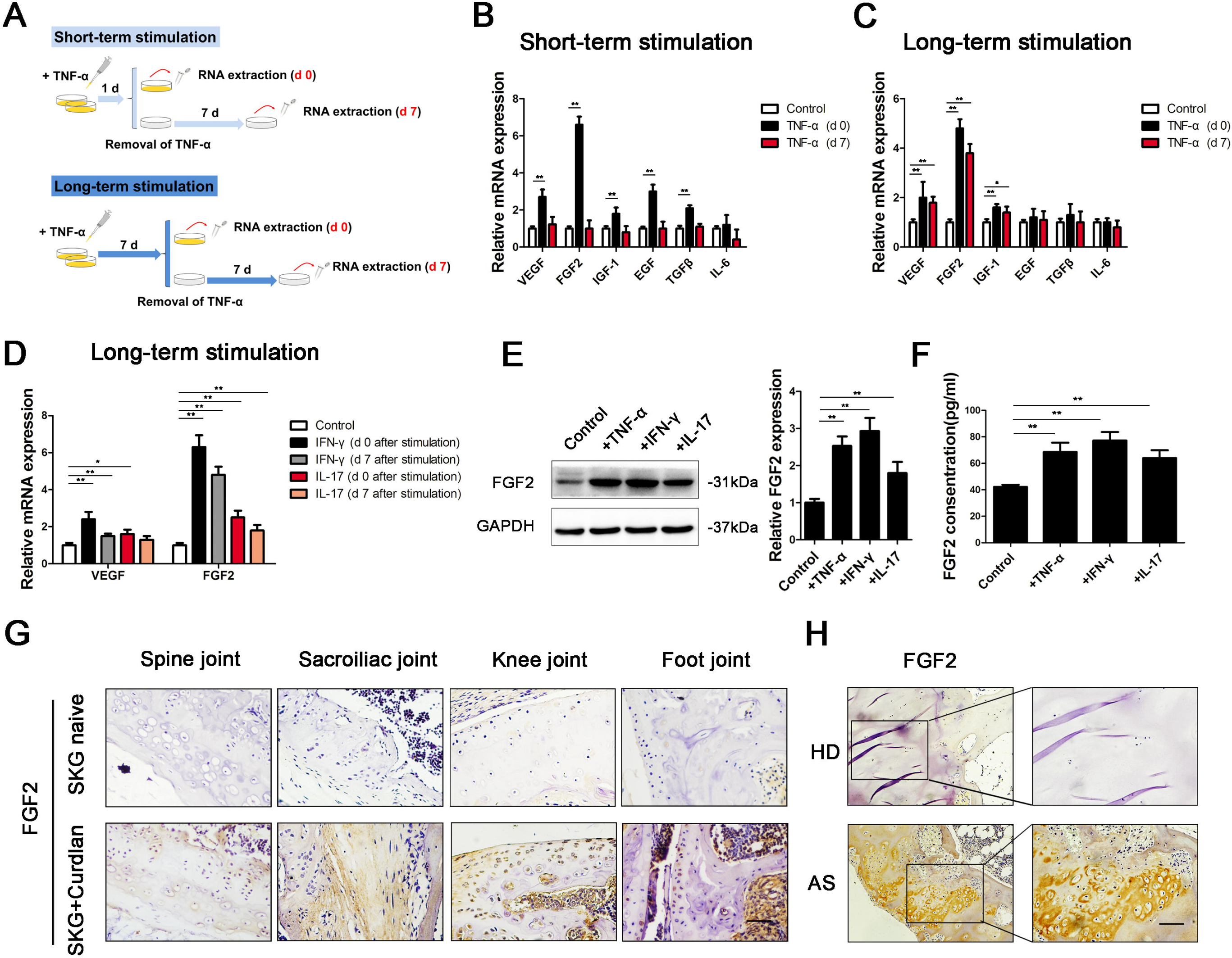
Long-term stimulation with inflammatory factors increased the expression of FGF2 in chondrocytes. (a) Diagram of 2 different modes of TNF-α stimulation in chondrocytes. (b&c) The expression of angiogenic factors (VEGF, FGF2, IGF-1, EGF, TGF-β, and IL-6) was measured in chondrocytes cultured with TNF-α for 1 day (b) or 7 days (c). (d) qRT-PCR analysis of VEGF and FGF2 expression in chondrocytes cultured with IFN-γ or IL-17 for 7 days, followed by removal of the stimulus for 0 days or 7 days, respectively. (e&f) FGF2 expression was measured by WB (d) and ELISA (e) in chondrocytes cultured with TNF-α, IFN-γ or IL-17 for 7 days. (g) IHC analysis of the spine joint, sacroiliac joint, knee joint and foot joint of SKG mice. Scale bar, 50 μm. (h) IHC analysis of the hip joint of AS patients. Scale bar, 100 μm. Values are shown as the mean ± SD from 1 representative experiment of 3 independent experiments each performed in triplicate. *P < 0.05, **P < 0.01.

Previous studies showed that curdlan-treated SKG mice had arthritis not only in peripheral joints but also in axial joints.^23,24^ Eight weeks after treatment with curdlan, SKG mice exhibited signs of inflammation (Supplementary Fig. 2). We then sacrificed the animals and performed IHC staining for FGF2 in the axial joints (spine, sacroiliac) and peripheral joints (knee, foot). The expression of FGF2 was higher in cartilage from curdlan-treated SKG mice than untreated SKG mice (Fig. 1g). Because cartilage is always replaced by bone in AS patients after surgery, it is difficult to obtain enough cartilage tissue for chondrocyte extraction from AS patients. Alternatively, in AS patients, we performed hematoxylin and eosin (H&E) and IHC staining on the femur head, which contains a small amount of residual cartilage. The results showed that the remaining cartilage was invaded by granulation tissue and expressed higher levels of FGF2 than the femur head in non-AS individuals (Fig. 1h, Supplementary Fig. 3). These results suggest that chronic inflammation induces FGF2 expression in chondrocytes in AS patients.

### Chondrocytes secrete FGF2 during chronic inflammation to induce angiogenesis

To explore whether FGF2 is an important factor for angiogenesis during the long-term stimulation of chondrocytes, we performed tube formation and transwell migration assays. The results demonstrated that the angiogenic activities of HUVECs were enhanced by CM from chondrocytes following long-term stimulation with TNF-α, IFN-γ or IL-17. In contrast, treatment of HUVECs with FGF-2 NAb significantly abolished chondrocyte CM-induced HUVEC tube formation and migration (Fig. 2a-d). To further characterize the angiogenic function of FGF-2 expression in chondrocytes, an *in vivo* Matrigel plug assay was performed. The results showed that plugs containing TNF-α-stimulated chondrocyte CM exhibited a notable increase in blood vessel growth, which was inhibited dramatically by FGF2 NAb (Fig. 2e-g). Therefore, proinflammatory factor-induced FGF-2 expression in chondrocytes subsequently promotes angiogenesis both *in vitro* and *in vivo*.

**Figure 2.**
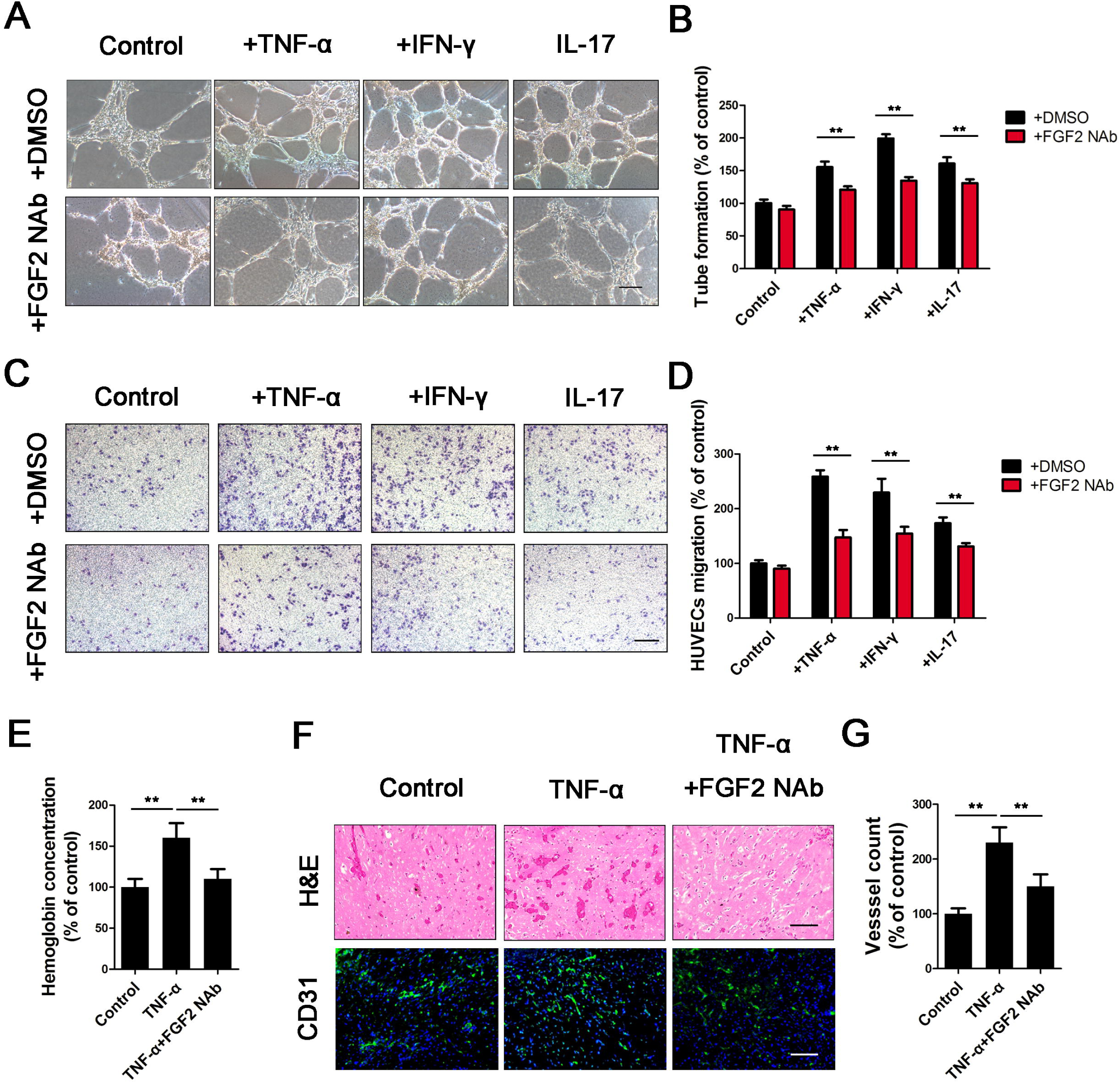
Chondrocytes secrete FGF2 to induce angiogenesis. The conditioned media (CM) from chondrocytes cultured with TNF-α, IFN-γ or IL-17 for 7 days were collected separately and then applied to HUVECs. (a-d) Tube formation (a) and transwell migration (c) assays were performed on HUVECs with CM. The numbers of branches (b) and migratory cells (d) were calculated and quantified using ImageJ software. Scale bar, 100 μm. (e-g) Matrigel plugs containing chondrocyte CM, TNF-α-treated chondrocyte CM or TNF-α-treated chondrocyte CM with FGF-2 NAb were subcutaneously injected into nude mice. The plugs were collected on day 7. Hemoglobin levels in the plugs were quantified and normalized to that in the control group (e). Paraffin sections of Matrigel plugs were stained with H&E and CD31 (f). Scale bar, 100 μm, n = 6. The numbers of vessels were quantified (g). Values are shown as the mean ± SD from 1 representative experiment of 3 independent experiments each performed in triplicate. *P < 0.05, **P < 0.01.

### Chronic inflammation triggers ERS and enhances the angiogenic ability of chondrocytes

Because inflammation is an important cause of ERS,^25^ we examined the expression of ERS-related genes in chondrocytes. The expression of GRP78 and GRP94 in chondrocytes was significantly increased after 7 days of stimulation with inflammatory factors (Fig. 3a&c). The unfolded protein response (UPR) is the major protective mechanism by which ERS is relieved and involves 3 pathways (IRE1a, PERK and ATF6).^18^ Further detection of IRE1a, PERK and ATF6 revealed that their expression in stimulated chondrocytes was higher than that in control chondrocytes (Fig. 3b). The expression of ATF6 was highest among proteins involved in the 3 key pathways (Fig. 3b&c). We also examined chondrocyte ERS levels *in vivo* by IHC staining on cartilage samples collected from AS patients. Both GRP78 and ATF6 were highly expressed in the chondrocytes of AS patients (Fig. 3d).

**Figure 3.**
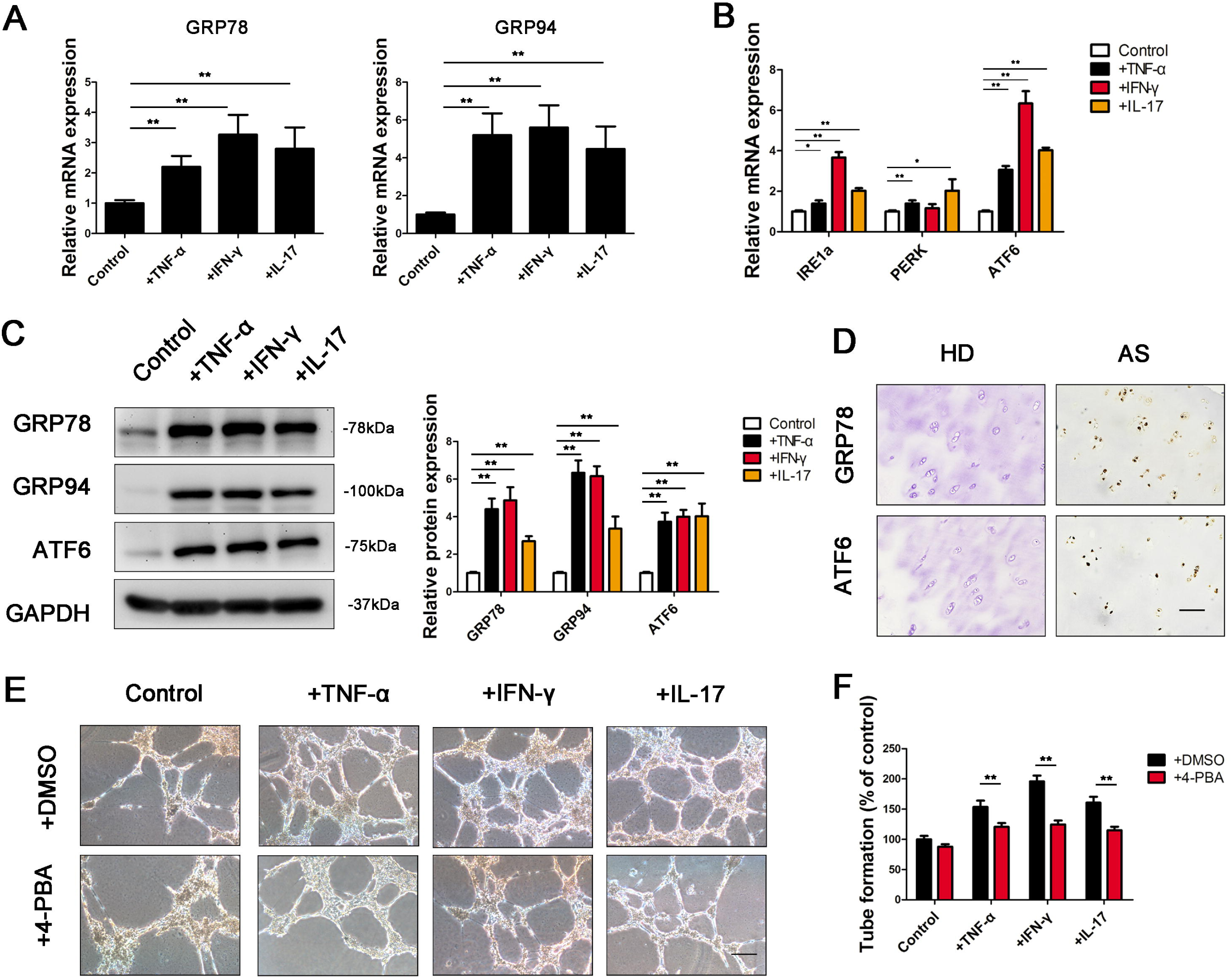
Long-term stimulation with inflammatory factors triggers ERS in chondrocytes and enhances angiogenesis. (a&b) The expression of ERS genes (GRP78, GRP94; a) and UPR pathway genes (IRE1a, PERK, and ATF6; b) was measured in chondrocytes cultured with TNF-α, IFN-γ or IL-17 for 7 days. (c) Protein levels of GRP78, GRP94, and ATF6 were measured with WB and quantified using ImageJ software. (d) IHC analysis of GRP78 and ATF6 expression in the hip joint of AS patients. Scale bar, 100 μm. (e&f) Tube formation assays were performed on HUVECs with chondrocyte CM (e). The numbers of branches were calculated and quantified using ImageJ software (f). Scale bar, 100 μm. Values are shown as the mean ± SD from 1 representative experiment of 3 independent experiments each performed in triplicate. *P < 0.05, **P < 0.01.

Previous studies have shown that ERS induces angiogenic activity.^16^ We thus explored whether the angiogenic effect of stimulated chondrocytes was related to ERS. The ERS alleviator 4-PBA significantly reduced the ability of stimulated chondrocytes to promote tube formation and migration of HUVECs (Fig. 3e&f, Supplementary Fig. 4). All these results indicate that proinflammatory factors in AS stimulate chondrocytes and promote angiogenic activity by increasing ERS levels.

### ERS induces FGF2 expression in chondrocytes through the ATF6 pathway

While many studies have focused on the relation between IRE1a or PERK and angiogenesis, few have proven the angiogenic role of ATF6. As the expression of ATF6 was the highest in proinflammatory factor-stimulated chondrocytes, we explored the role of ATF6 in the angiogenic effect of stimulated chondrocytes. First, chondrocytes were transfected with plasmids encoding an shRNA targeting ATF6 (Supplementary Fig. 5). We then examined FGF2 and GRP78 (a known target gene of ATF6) expression in control and proinflammatory factor-stimulated chondrocytes. FGF2 and GRP78 expression was significantly reduced by ATF6 knockdown in stimulated chondrocytes but remained unchanged in control chondrocytes (Fig. 4a&b). Furthermore, tube formation and transwell migration assays showed that the knockdown of ATF6 significantly reduced the angiogenic effect of stimulated chondrocytes but not control chondrocytes (Fig. 4c, Supplementary Fig. 6).

**Figure 4.**
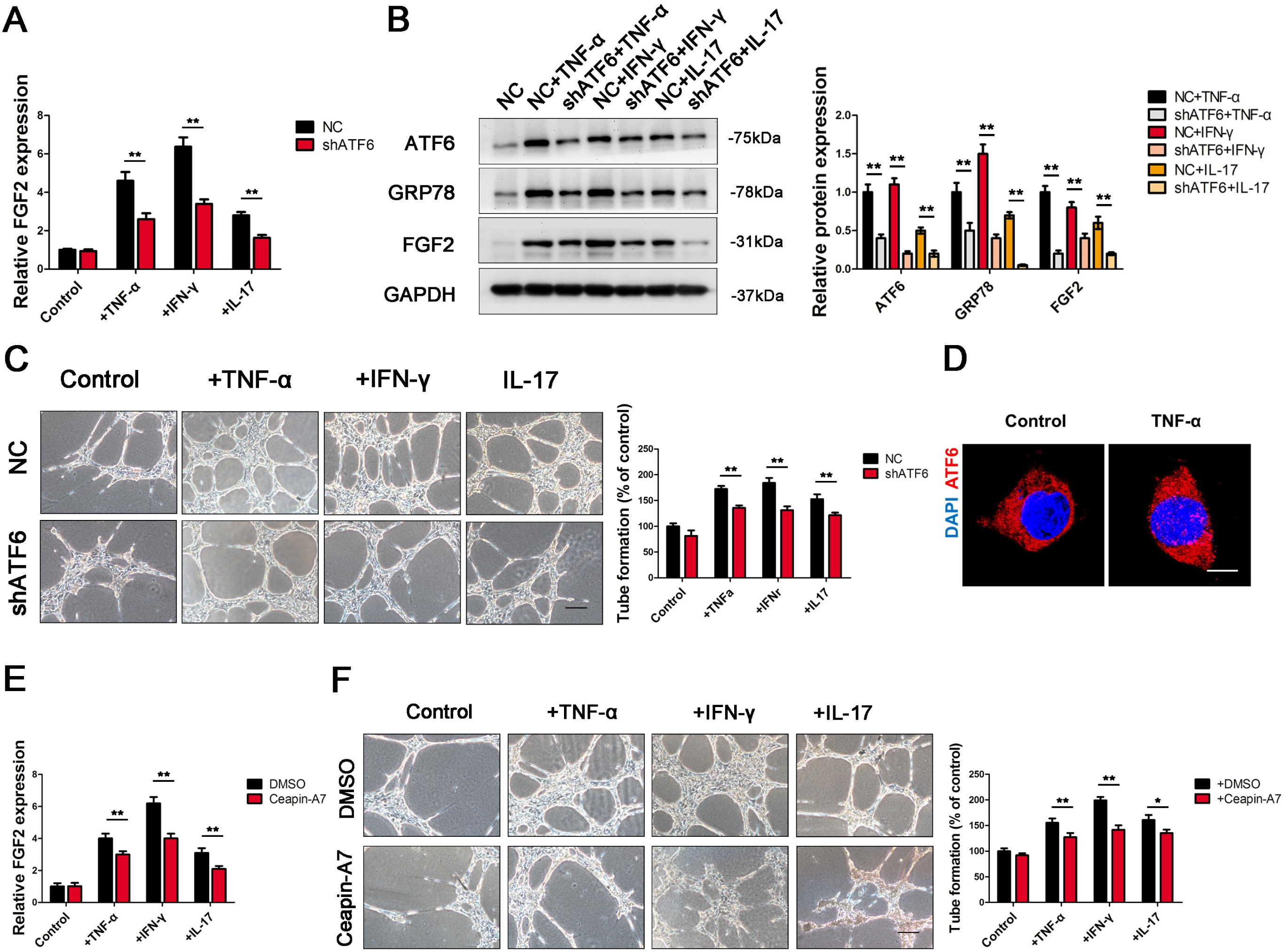
Activation of the ATF6 pathway promoted FGF2 expression in chondrocytes. (a) FGF2 mRNA levels were measured in chondrocytes with ATF6 knockdown. (b) Protein levels of ATF6, GRP78, and FGF2 were measured with WB and quantified using ImageJ software. (c) Tube formation assays were performed. The numbers of branches were calculated and quantified using ImageJ software. Scale bar, 100 μm. (d) Confocal imaging showed that ATF6 translocated to the nucleus in chondrocytes after TNF-α stimulation. (e) qRT-PCR analysis of FGF2 expression in chondrocytes treated with or without Ceapin-A7. (f) Tube formation assays were performed with CM derived from chondrocytes treated with or without the ATF6 activation inhibitor Ceapin-A7. The numbers of branches were calculated and quantified using ImageJ software. Scale bar, 100 μm. Values are shown as the mean ± SD from 1 representative experiment of 3 independent experiments each performed in triplicate. *P < 0.05, **P < 0.01.

ATF6 activation includes ATF6 cleavage by S1P and S2P in the Golgi apparatus and thus translocation to the nucleus.^20^ An analysis of confocal images showed that ATF6 translocated to the nucleus of chondrocytes after stimulation with TNF-α (Fig. 4d). Ceapin-A7 is a specific inhibitor of ATF6 activation that functions by inhibiting the transport of ATF6 from the ER to the Golgi apparatus.^26^ To explore whether FGF2 expression and the angiogenic effect of stimulated chondrocytes were related to activated or inactivated ATF6, we treated stimulated chondrocytes with Ceapin-A7. The inhibition of ATF6 activation by Ceapin-A7 decreased FGF2 expression (Fig. 4e, Supplementary Fig. 7a) and the angiogenic effect of stimulated chondrocytes (Fig. 4f, Supplementary Fig. 7b). Taken together, these results indicate that proinflammatory factors induce FGF2 expression in chondrocytes by activating the ATF6 pathway.

### ATF6 directly promotes FGF2 transcription

Activated ATF6 mediates gene expression by recognizing ERSE I (CCAAT-N9-CCAC(G)) and ERSE II (ATTGG-N1-CCAC(G)).^27^ To explore whether ATF6 promoted FGF2 expression in chondrocytes in a direct or an indirect manner, we scanned *FGF2* promoter regions spanning from −10,000 bp to 0 bp of the transcription start site (TSS) for a potential ERSE in various species. The analysis revealed *ERSE* sequences in the *FGF2* promoters of various species, including humans, mice, rabbits, cattle and sheep (Fig. 5a). We also identified 4 potential ERSEs in the promoters of *hFGF2*, among which 2 potential ERSEs, namely, M1 (−1420 to −1402) and M2 (−1784 to −1766), were located at −2000-0 bp (Fig. 5b). These results support that ATF6 directly binds to the FGF2 promoter and mediate its transcription.

**Figure 5.**
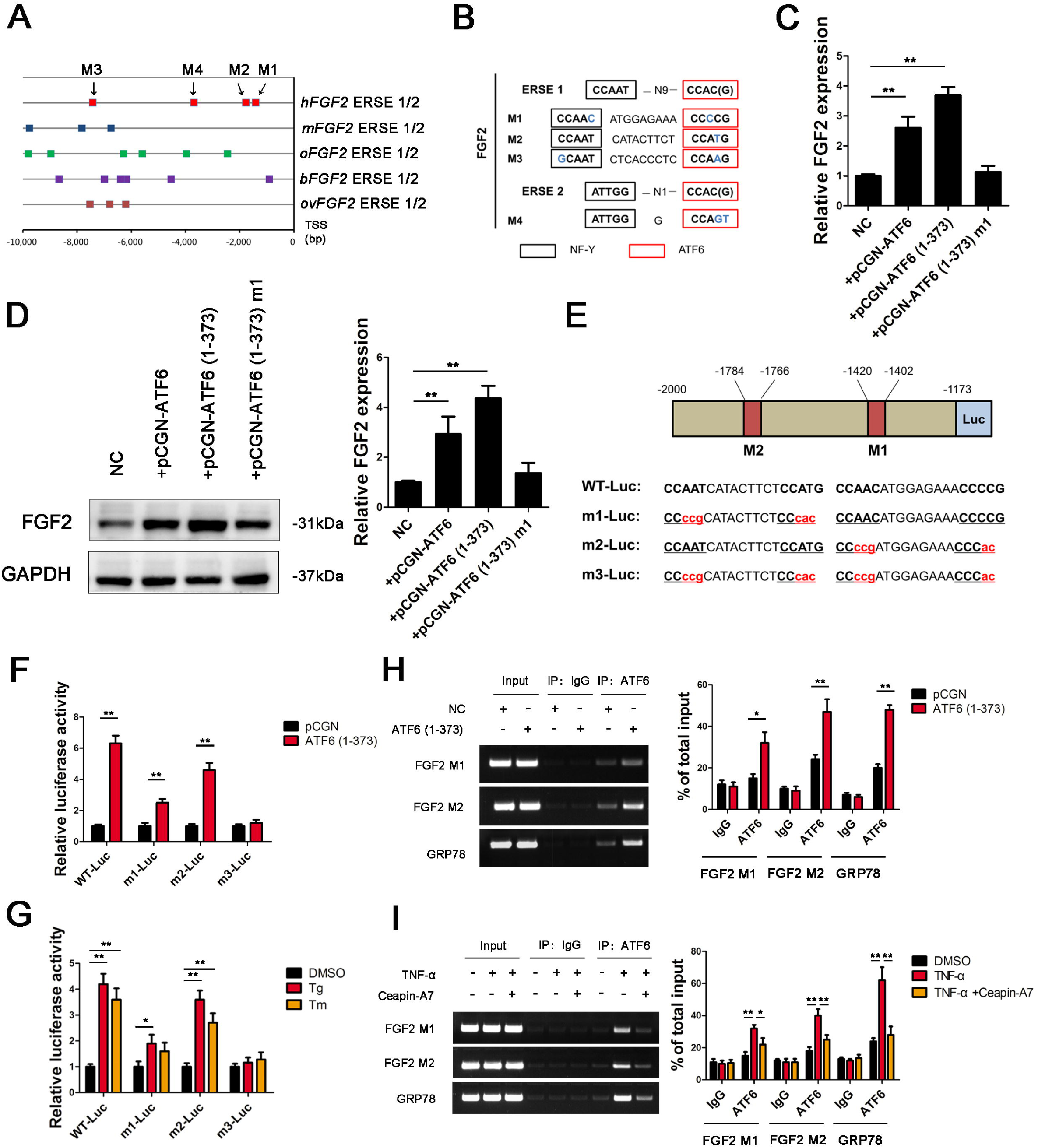
ATF6 directly binds to the *FGF2* promoter and promotes its transcription. (a) Schematic representation of the FGF2 promoters from human (*Homo sapiens, hFGF2*), mouse (*Mus musculus, mFGF2*), rabbit (*Oryctolagus cuniculus, oFGF2*), cattle (*Bos taurus, bFGF2*), and sheep (*Ovis aries, ovFGF2*). Different squares represent the possible ATF6-binding sites (ERSE I or ERSE II) of different species. TSS, transcriptional start site. (b) Diagram showing the noncanonical ERSE I and ERSE II in the *hFGF2* promoter. (c&d) Both mRNA (c) and protein (d) levels of FGF2 in chondrocytes were measured. (e) Diagram of the locations of ERSE I (M1, M2) in the *hFGF2* 5′-flanking region, their sequences (WT), and mutations to those sequences (m1, m2, and m3). (f&g) Chondrocytes were transfected with plasmids encoding *hFGF2*(−2000/-1173)-Luc WT, m1, m2, or m3. Chondrocytes were then transfected with ATF6-expressing vectors (f) or treated with ERS inducers (Tg and Tm; g). Forty-eight hours later, luciferase levels were measured in extracts. (h&i) After transfection with ATF6 (1-373) in chondrocytes for 48 h (h) or treatment with TNF-α for 7 days (i), the DNA-binding ability of ATF6 to the hFGF2 and GRP78 promoters was analyzed using ChIP assays. Values are shown as the mean ± SD from 1 representative experiment of 3 independent experiments each performed in triplicate. *P < 0.05, **P < 0.01.

To explore whether ATF6 directly binds to the FGF2 promoter and mediates transcription, we overexpressed full-length ATF6, the active form of ATF6 (1-373), or the negative mutation ATF6 (1-373) m1 in chondrocytes. FGF2 expression in chondrocytes was promoted by ATF6 and ATF6 (1-373) but not ATF6 (1-373) m1 (Fig. 5c&d). We then cloned a truncated version of the *hFGF2* 5’-flanking sequence (−2000 to −1173) in front of firefly luciferase (referred to as WT-Luc). M1 and M2 were mutated either singly or together (m1-Luc, m2-Luc, and m3-Luc) (Fig. 5e). As shown in Fig. 5f, transfection with ATF6 (1-373) significantly increased luciferase activity in WT-Luc-, m1-Luc- and m2-Luc-transfected 293T cells but not m3-Luc-transfected 293T cells, indicating that ATF6 binds to WT-Luc, m1-Luc and m2-Luc but not m3-Luc. We further used Tg and Tm to induce endogenous ATF6 expression. Tg and Tm significantly increased luciferase activity in WT-Luc-, m1-Luc- and m2-Luc-transfected 293T cells but not m3-Luc transfected 293T cells (Fig. 5g). Finally, we performed a ChIP assay to explore the DNA-binding ability of ATF6 to the FGF2 promoter. The results revealed that ATF6 bound to the M1 and M2 sequences in the FGF2 promoter, as well as the GRP78 gene promoters, in chondrocytes. The expression of not only exogenous active ATF6 but also endogenous ATF6 increased ATF6 binding to the FGF2 promoter, which was inhibited by Ceapin-A7 (Fig. 5h&i). These results suggest that ATF6 directly binds to the FGF2 promoter and promotes its transcription.

### ATF6 aggravates angiogenesis-osteogenesis coupling in mice

To observe the effect of ERS on angiogenesis and osteogenesis, we performed an intra-articular injection of Tg to induce ERS in mouse chondrocytes (Supplementary Fig. 8a). IHC staining was also performed on the knee joints of experimental mice. Tg treatment significantly increased the expression of ATF6, GRP78 and FGF2. However, after treatment with the ATF6 inhibitor Ceapin-A7, the expression of ATF6 was not changed, while the expression of GRP78 and FGF2 was significantly decreased (Supplementary Fig. 8b), which suggested that FGF2 and GRP78 act downstream of ATF6 in mouse chondrocytes. To explore the changes in angiogenesis and osteogenic activity in the knee joint, we performed IF staining of the endothelial cell marker CD31 and the osteoblast marker OCN in experimental mice. As shown in Supplementary Fig. 8c&d, Tg treatment significantly increased CD31^+^ vessels and OCN^+^ cells near the endosteum in bone marrow, suggesting that ERS leads to increased angiogenesis and osteogenic activity. However, after treatment with the ATF6 inhibitor Ceapin-A7, CD31^+^ vessels and OSX^+^ cells were significantly reduced, suggesting that the enhancement of angiogenesis and osteogenic activity is achieved mainly through the ATF6 pathway.

We further treated SKG mice with Ceapin-A7. One week after curdlan induction, SKG mice were treated with drinking water containing DMSO or 1 μM Ceapin-A7. Because a previous study showed that ATF6 antagonized chronic liver injury,^20,28^ we tested alanine aminotransferase (ALT) and aspartate aminotransferase (AST) levels in the sera of SKG mice. Neither ALT nor AST levels were elevated in Ceapin-A7-treated mice (Fig. 6a), indicating that Ceapin-A7 at such does not cause liver damage in mice. To determine whether Ceapin-A7 influences the inflammatory state in SKG mice, we also examined the local expression of TNF-α, IFN-γ and IL-17 in hind paws (Fig. 6b). TNF-α, IFN-γ and IL-17 levels did not change in Ceapin-A7-treated mice, indicating that Ceapin-A7 does not influence the inflammatory state in SKG mice. Twelve weeks after curdlan induction, SKG mice developed severe arthritis. IF staining of CD31 and OCN in the vertebral body showed that the numbers of CD31^+^ vessels and OCN^+^ cells were increased in untreated SKG mice compared with Ceapin-A7-treated SKG mice (Fig. 6c&d). We also performed a micro-CT scan and found bony bridging between vertebrae (spinal ankylosis) (6/10, n=10). Following treatment with Ceapin-A7, the occurrence of spinal ankylosis decreased (1/10, n=10) (Fig. 6e). Similar osteogenic changes were observed in the hind paws. New bone formed in the hind paws (16/20, n=10). Following treatment with Ceapin-A7, the incidence of osteophyte formation was significantly reduced (4/20, n=10) (Fig. 6f). Moreover, IHC staining showed low FGF2 expression in the cartilage of SKG mice treated with Ceapin-A7 (Fig. 6e&f). These results indicate that the ATF6 inhibitor Ceapin-A7 slows the pathological progression of osteogenesis by inhibiting angiogenesis-osteogenesis coupling. Taken together, these data suggest that chondrocytes stimulated by chronic inflammation undergo chronic ERS via activation of the ATF6 pathway and thus secrete more FGF2 to aggravate angiogenesis-osteogenesis coupling in AS (Fig. 7).

**Figure 6.**
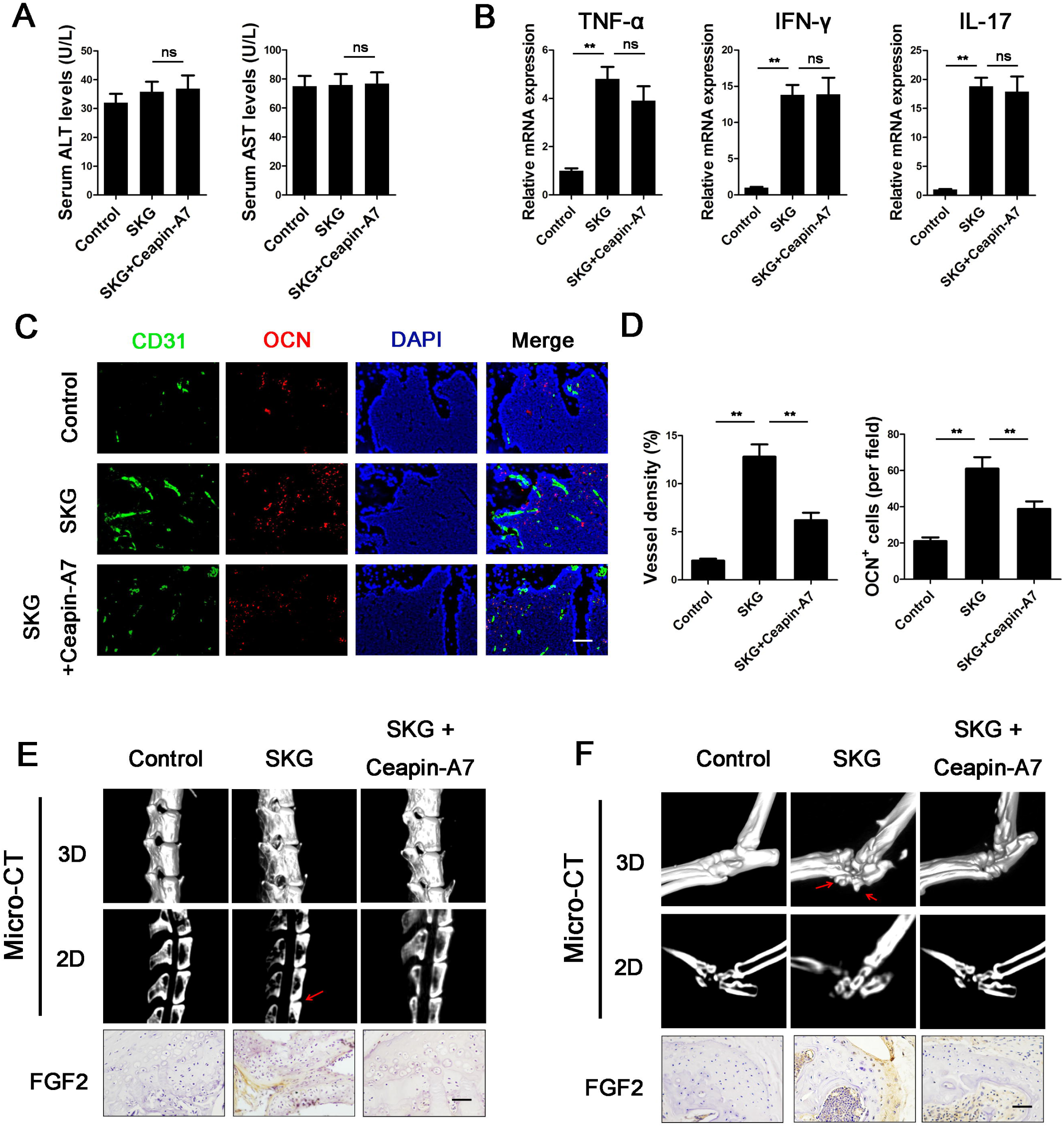
ATF6 regulates angiogenesis-osteogenesis coupling through FGF2 *in vivo*. (a) ALT and AST levels in the serum of SKG mice were measured. (b) Local mRNA expression of TNF-α, IFN-γ and IL-17 in the hind paws of SKG mice. (c) IF staining of vertebral bodies from SKG mice showing endothelial cells (CD31^+^) and osteoblasts (OCN^+^). Scale bar, 50 μm. (d) Quantification of the proportions of CD31^+^ vessels and OCN^+^ cells. (e&f) Representative micro-CT images of a spine (e) and ankle (f) obtained from SKG mice at baseline, 12 weeks after induction, and 12 weeks after induction and treatment with Ceapin-A7. IHC staining of FGF2 in the spine and ankle. Values are shown as the mean ± SD. *P < 0.05, **P < 0.01, ns, not significant.

**Figure 7.**
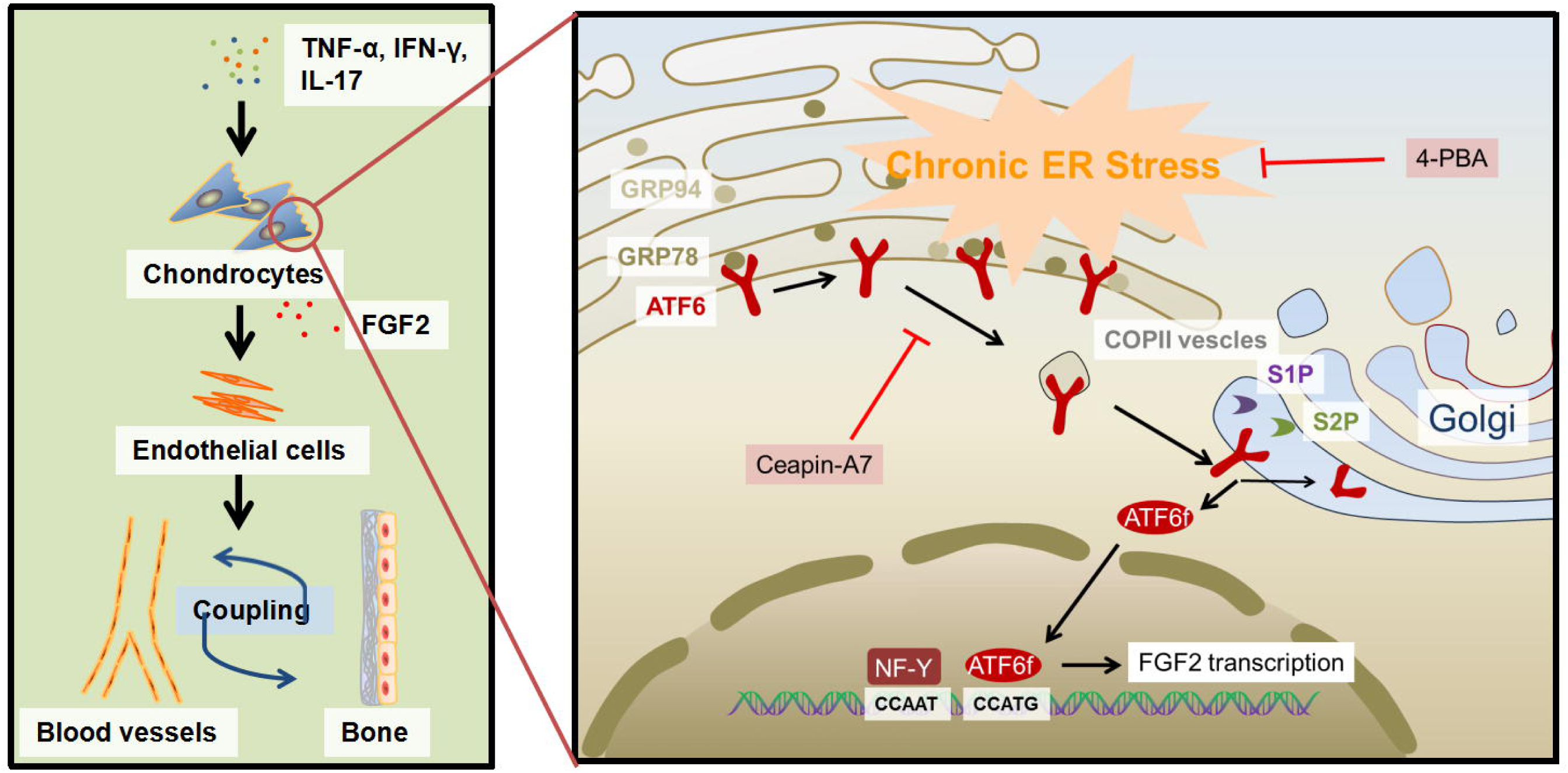
Proposed model of the relationship between ATF6 and FGF2 in chondrocytes in AS. FGF2 is an effective factor that promotes angiogenesis-osteogenesis coupling and plays an important role in AS. Chondrocytes stimulated long-term with proinflammatory cytokines undergo chronic ERS with ATF6 pathway activation. By directly binding to the *FGF2* promoter, ATF6 increases FGF2 expression. Treatment of chondrocytes with the ATF6 inhibitor Ceapin-A7 effectively inhibited FGF2 expression.

## Discussion

Though inflammation is an important pathologic process in AS, the association between inflammation and pathologic bone formation has not yet been illuminated. More than 10 years ago, some researchers hypothesized that enthesitis was uncoupled with pathologic bone formation in AS patients.^6^ *In vitro* experiments showed that proinflammatory factors such as TNF-α and IL-1 accelerated the activity of osteoclasts while inhibiting the differentiation of osteoblasts.^29^ Furthermore, many mechanisms other than inflammation, such as BMP, Wnt, Hh signaling and mechanical loading can explain pathologic bone formation in AS.^30–32^ We previously showed that mesenchymal stem cells from AS patients secreted more BMP-2 but less Noggin coupled with abnormal osteogenic differentiation than cells from healthy individuals.^33^ However, whether these signaling pathways act downstream of inflammation or independent of inflammation remains unknown. In our study, we showed that proinflammatory factor stimulation promoted the prolonged angiogenesis effect of chondrocytes, even after removal of the proinflammatory factors. This result indicates that we should take into account the interval between two inflammatory periods during which there might be abnormal osteogenic activity.

The structural destruction and replacement of cartilage by granulation tissue is an important pathological change observed in AS patients.^11^ Actually, an abnormal cartilage structure might be a common change before osteophyte formation. In 2009, Benjamin et al.^34^ applied IHC assay to spine osteophytes from the elderly and found a mixture of bone, cartilage, blood vessels and other tissues. They also observed active osteogenesis and osteolysis activity among osteophytes. This discovery indicated that the formation of osteophytes is the result of cartilage calcification, during which blood vessels play an important role. Additionally, IHC staining on the hip joints and facet joints from AS patients revealed an abnormal cartilage structure, inside of which blood vessels and granulation tissue, accompanied by active osteogenesis and osteolysis activity, were observed.^14,15^ Taken together, these results suggest that blood vessel invasion is closed related to cartilage destruction and calcification.^11^ However, the mechanism of blood vessel invasion remains unknown. As we showed above, chondrocytes treated with proinflammatory factors secrete a variety of angiogenic factors and promote endothelial cell migration and lumen formation. Similarly, in hip joints from AS patients, chondrocytes express high levels of FGF2. Therefore, we hypothesize that inflammation is a major cause of the structural destruction of cartilage by promoting vascular invasion. Further study is needed to clarify the role of inflammation in the structural destruction of cartilage and pathologic bone formation.

Currently, whether ERS plays a role in AS development remains controversial.^35^ Although several lines of evidence have demonstrated high ERS in HLA-B27 transgenic animals, data from AS patients are lacking. Dong et al.^36^ revealed that macrophages in peripheral joints from AS patients expressed high levels of GRP78, which indicated high levels of ERS. However, some studies showed that the ERS level in several tissues, including the peripheral blood, synovia and intestine, from AS patients did not differ from that in controls.^37^ We found that the levels of GRP78 and ATF6 were higher in chondrocytes in the femur heads of AS patients than in those of controls, prompting chondrocytes in AS patients to have high levels of ERS. Similarly, high levels of GRP78 and ATF6 were found in the chondrocytes of SKG mice. In addition, we treated chondrocytes with proinflammatory factors for 7 days and successfully prolonged ERS. All these results indicate that certain cell types in AS patients have a high level of ERS and might be induced by chronic inflammation.

ERS is a key regulator of angiogenesis.^16^ Many studies have focused on the mechanisms by which ERS regulates angiogenic activities. A previous study showed that the expression of the key angiogenic factor VEGF was under the control of Xbp-1, ATF4 and ATF6.^38^ Ghosh et al.^19^ found that ATF6 promoted VEGF transcription by directly binding to its gene promoter in the liver cancer cell line HepG2. However, in our study, we did not observe a change in VEGF mRNA levels in chondrocytes after ATF6 knockdown. This result suggested that ATF6 was not a key transcription factor by which VEGF transcription was regulated in chondrocytes. Moreover, we noted that FGF2 mRNA levels decreased after ATF6 knockdown or ATF6 inhibition in chondrocytes. We also proved that ATF6 directly binds to the *FGF2* promoter region in human chondrocytes. These results indicate that ATF6-mediated FGF2 transcription is an important pathway by which ERS regulates angiogenic activities.

FGF2 is a well-studied polypeptide that functions in angiogenesis, tissue regeneration and neuroprotection.^39–41^ Several lines of evidence suggest that FGF2 promotes cartilage repair and bone growth^42–44^ and plays an important role in the process of osteoarthritis.^45,46^ However, the direct regulatory effect of FGF2 on osteoblasts is still under debate. Some studies have suggested that the FGF2 signaling pathway enhances the transcriptional activity of Runx2 and promotes osteoblast differentiation,^47^ while others have indicated that FGF2 inhibits osteoblast differentiation.^48^ Nevertheless, it is hypothesized that the effect of FGF2 on cartilage repair and bone growth depends on its angiogenesis promotion effect.^44^ By performing IHC staining on sections of femoral heads from AS patients, we found that FGF2 was highly expressed in femoral head chondrocytes of AS patients and an increase in vascular structures in cartilage tissues. Based on the results described above, we conclude that FGF2 plays an important role in the processes of cartilage calcification and pathologic osteogenesis, and it is likely to be achieved by promoting vascular growth.

In summary, we identified ATF6 as a critical positive regulator of angiogenesis-osteogenesis coupling in AS. Activated by long-term inflammation, ATF6 increases FGF2 expression by mediating FGF2 transcription in chondrocytes. This study also demonstrated that ATF6 in chondrocytes may represent a promising therapeutic target for AS.

## Methods

### Patient recruitment

This study was approved by the Ethics Committee of The Eighth Affiliated Hospital, Sun Yat-sen University, Shenzhen, China. Twenty trauma patients and twenty AS patients who underwent hip replacement surgery were recruited, and femur heads were collected. All the AS patients met the modified New York criteria. Written informed consent was obtained from all participants.

### Cell culture and treatment

Primary human articular chondrocytes were collected from the articular cartilage through enzymatic digestion. Briefly, cartilage slices were minced and washed clean with 1% PBS, followed by digestion in a mixture of 0.25% collagenase II (Gibco) in FBS (Hangzhou Sijiqing Biological Engineering Material Co. Ltd., China) free DMEM (glucose 4500 mg/L; Gibco) with Pen-Strep (Guangzhou Jingxin Biotechnology Co., Ltd., China) at 37°C for 3-5Lh in a spinner flask in an incubator with a 5% CO_2_ atmosphere. Then, the cell suspension was used to establish cultures in T75 flask. At 3-4 days after harvesting, primary chondrocytes were re-plated at 80% to 85% confluence in 6-well plates and used for further studies. Additionally, the chondrocytes were validated through the immunofluorescence staining of collagen II.

Human umbilical vein endothelial cells (HUVECs) and the human embryonic kidney cell line (293T) were obtained from iCell Bioscience (Shanghai, China). HUVECs were cultured in endothelial growth medium 2 (EGM-2; Lonza, Switzerland) containing 10% FBS and the supplied growth factors, and cells at the third to fourth passages were used in all experiments. 293T cells were cultured in high-glucose DMEM supplemented with 10% FBS.

The ERS inducers tunicamycin (Tm, 1 μg/ml) and thapsigargin (Tg, 1 μM), the ERS alleviator 4-phenyl butyric acid (4-PBA, 5 mM) and the ATF6 inhibitor Ceapin-A7 (500 nM) were purchased from Sigma-Aldrich. Chondrocytes were cultured in DMEM supplemented with Tm or Tg for 6 h. Chondrocytes were cultured in DMEM supplemented with 4-PBA or Ceapin-A7 for 24 h. The inflammatory cytokines TNF-α (10 ng/ml), IFN-γ (500 U/L) and IL-17 (10 ng/ml) were obtained from PeproTech. For short-term stimulation, chondrocytes incubated with TNF-α were treated for 1 day. For long-term stimulation, chondrocytes incubated with TNF-α, IFN-γ or IL-17 were cultured for 7 days, and the medium was changed every other day. Then, the stimulus was removed, and the samples were cultured in DMEM and harvested after an additional 7 days. After the stimulus was removed, the samples were cultured in EGM-2 for 24 h to produce conditioned media (CM) for further experiments (Fig. 1a).

### Mice

All animal care protocols and experiments were reviewed and approved by the Ethics Committee of Sun Yat-sen University. C57BL/6N male mice and BALB/c male nude mice (4 weeks old) were purchased from the Guangdong Medical Laboratory Animal Center (Guangdong, China). SKG mice were obtained from S. Sakaguchi (University of Kyoto, Kyoto, Japan) and bred in house. Disease was induced between 6 and 10 weeks of age using 3 mg curdlan (InvivoGen) administered by intraperitoneal injection. Clinical features in the mice were monitored weekly and scored by the same observer, who was blinded with regard to treatment, as follows: 0, no swelling or redness; 0.1, swelling or redness of digits; 0.5, mild swelling and/or redness of wrists or ankle joints; and 1, severe swelling of the larger joints. The scores of the affected joints were summed; the maximum possible score was 6. All mice were housed in a specific pathogen-free facility.

### Plasmid transfection

Plasmid transfections were performed using Lipofectamine 3000 (Invitrogen) according to the manufacturer’s instructions. The siRNAs and siATF6 were purchased from GenePharma (Shanghai, China). pCGN-ATF6, pCGN-ATF6 (1-373) and pCGN-ATF6 (1-373) m1 were gifts from Professor Ron Prywes (Addgene plasmid 11974, 27173, 27174).

### Quantitative real-time PCR (qRT-PCR)

Total RNA was extracted from cultured cells or tissues using TRIzol Reagent (Invitrogen) and transcribed into cDNAs using a PrimeScriptTM RT Reagent Kit (Takara Bio, Mountain View, CA). qRT-qPCR was performed with a SYBR PrimeScript RT-qPCR Kit (Takara) for 40 cycles at 95°C for 5 s and 60°C for 30 s on a Light Cycler 480 Real-Time PCR System (Roche, Switzerland). All primer sequences used in this study are listed in Supplementary materials (Supplementary Table S1).

### Western blotting (WB)

Cells were harvested and lysed in RIPA buffer containing a protease inhibitor cocktail (1:100). Total protein was extracted, and the concentration was determined by the BCA method (Thermo Fisher). Then, equal amounts of total protein were separated by 10% SDS-PAGE and transferred onto polyvinylidene difluoride (PVDF) membranes (Millipore). The membranes were blocked and incubated with primary antibodies against FGF2 (36769, SAB), ATF6 (32008, SAB), GRP78 (3177S, CST), GRP94 (20292, CST), and GAPDH (5174, CST). Protein bands were then incubated with secondary anti-mouse or anti-rabbit peroxidase-linked antibodies (Beyotime). We then used enhanced chemiluminescent (ECL) detection reagents (Beyotime) to obtain the final result. Three replicates were performed for all WB analyses. WB results were quantified using ImageJ software (National Institutes of Health, Bethesda, MD).

### Enzyme-linked immunosorbent assay (ELISA)

To measure the FGF-2 production of chondrocytes in the CM, the cells were seeded into 6-well plates (6×10^4^ cells/well). Then, the cells were treated with TNF-α (10 ng/ml), IFN-γ (500 U/L) and IL-17 (10 ng/ml) and incubated in a humidified incubator at 37°C for 7 days. After incubation, the supernatant CM was collected and stored at −80L until the assay was performed. FGF-2 in the CM was assayed using a human FGF basic DuoSet ELISA Development Kit (R&D Systems) according to the manufacturer’s procedure.

### Immunohistochemistry (IHC) and immunofluorescence (IF)

Paraffin-embedded sections were prepared, mounted onto slides, deparaffinized in xylene, rehydrated in a graded alcohol series and washed in deionized water. After antigen retrieval (sections were heated at 95-100 □ on a hotplate for 30 min in 10 mM sodium citrate, pH 6.0), intrinsic peroxidase activity was blocked by incubation with 3% H_2_O_2_. Nonspecific antibody-binding sites were blocked using 3% BSA in PBS. Sections were then incubated with appropriately diluted primary antibodies specific for human or mouse CD31 (3568, Abcam), osteocalcin (OCN; ab93876, Abcam), FGF2 (36769, SAB), ATF6 (32008, SAB) or GRP78 (3177S, CST) at 4°C overnight. Then, we followed the instructions of the SP Rabbit & Mouse HRP Kit (CWBio) for IHC. Alternatively, we incubated the slides with a fluorescence-labeled secondary antibody (CST). Slides were observed under a light microscope or a fluorescence microscope.

### Transwell migration assay

The transwell migration assay was performed using Transwell inserts (8.0 μm pore size; Costar) in 24-well plates. HUVECs (10^4^ cells in 200 μl of EGM-2with 10% FBS) were then seeded into the upper chamber, and 300 μl of chondrocyte CM was placed in the lower chamber. Cells on the lower side of the Transwell membrane were examined and counted under a microscope after crystal violet staining.

### Tube formation assay

Matrigel (BD Biosciences) was melted at 4°C, added to angiogenesis μ-slides (81506, Ibidi, Germany) at 10 μl/well, and then incubated at 37°C for 30 min. HUVECs (2×10^3^ cells) were resuspended in a 1:1 mixture of EGM-2 serum-free medium and chondrocyte CM (total 50 μl) and added to the wells. After 12 h of incubation at 37°C, HUVEC tube formation was assessed by microscopy, and each well was photographed under a light microscope. The numbers of branches were calculated and quantified using ImageJ software.

### In vivo Matrigel plug assay

Four-week-old male nude mice were divided into three groups (n = 10 for each group) and subcutaneously injected with 150 ul of Matrigel containing chondrocyte CM, respectively, control CM, TNF-α-treated CM, and TNF-α-treated CM with FGF-2 neutralizing antibody (NAb; AF-233, R&D Systems). On day 7, the Matrigel plugs were harvested: some were fixed with 3.7% paraformaldehyde for at least 2 days and then embedded in paraffin and subsequently processed for IHC and IF staining, whereas others were evaluated by Drabkin’s method (Drabkin’s Reagent Kit, Sigma-Aldrich) to quantify the hemoglobin concentration.

### Chromatin immunoprecipitation (ChIP) assay

ChIP assays were performed according to the manufacturer’s instructions provided in the SimpleChIP Enzymatic ChIP Kit (CST). Briefly, the ChIP assay was performed using protein A/G agarose (Thermo Fisher Scientific) and an anti-ATF6 antibody (ab227830, Abcam). The immunoprecipitated DNA was used to amplify DNA fragments via PCR with specific primers. The primer sequences for GRP78 were 5’-GCG GAG CAG TGA CGT TTA TT-3’ (forward) and 5’-GAC CTC ACC GTC GCC TAC T-3’ (reverse). The primer sequences for FGF2 M1 were 5’-TCTGAGCAAATAGGCCTTGCT-3’ (forward) and 5’-GGCTGAAGCCCCTGTAACAAA-3’ (reverse). The primer sequences for FGF2 M2 were 5’-ATATGCCTGGTTTTGGGCCT-3’ (forward) and 5’-CAGCCTACCGAATAGCTGGG-3’ (reverse).

### Construction of reporter plasmids and luciferase assays

FGF2-ERSE (ERSE-like site) was cloned into the pGL3-basic luciferase reporter plasmid. Chondrocytes (2.5×10^4^ cells per well) were seeded in triplicate into 24-well plates (Corning). After incubation for 24 h, the cells with either ATF6 plasmids or ATF6-mutant plasmids were transfected with 200 ng of FGF2-ERSE-luciferase-reporter plasmids using Lipofectamine 3000 (Invitrogen) according to the manufacturer’s recommendations. Each transfection included the same amount of Renilla, which was used to standardize transfection efficiency. Then, the cells were allowed to recover in medium containing 10% FBS for 24 h. Forty-eight hours post-transfection, firefly and Renilla signals were measured using a Dual Luciferase Reporter Assay Kit (Promega) and are presented as the increase in activation over the reporter alone.

### Micro-CT analysis

Lumbar spine (spinal segment including intervertebral disc and adjacent end plates) and hind paw specimens were obtained from mice postmortem and fixed with 4% paraformaldehyde. For micro-CT scans, specimens were fitted in a cylindrical sample holder and scanned using a Scanco lCT40 scanner set to 55 kVp and 70 lA. For visualization, the segmented data were imported and reconstructed as 3-dimensional images using MicroCT Ray software V3.0 (Scanco Medical).

### Statistical analysis

Statistical analyses were performed with SPSS 13.0 (SPSS, Inc., Chicago, IL, USA). Data are shown as the mean ± SD. For comparisons of 2 groups, a 2-tailed Student’s t-test was used. Comparisons of multiple groups were performed by using one-way analysis of variance (ANOVA) followed by Dunnett’s post hoc test. All experiments were repeated at least 3 times. Statistical significance was concluded at *P < 0.05 and **P < 0.01.

## Acknowledgements

This study was financially supported by grants from the National Natural Science Foundation of China (81971518), the National Natural Science Foundation of China (81871750), the Natural Science Foundation of Guangdong Province (2018A030313232), the Medical Science and Technology Research Project of Guangdong Province (A2018292), the Shenzhen Key Medical Discipline Construction Fund (ZDSYS20190902092851024) and the Health Welfare Fund Project of Futian District (FTWS2020078).

The authors thank Prof. Pheier Saw (Medical Research Center, Sun Yat-sen Memorial Hospital, SYSU, China) for her guidance.

## Declaration of interests

The authors declare no competing interests.

## Author contributions statement

MM, HL and PW planned the project, performed the experiments, interpreted the data and drafted the article. WY and RM designed the research and conducted data analysis. YJ and PS provided technical support. XS and YL collected the human samples and isolated and cultured the chondrocytes. YW and HS supervised the study.

## Supplementary material

**Supplementary Figure 1. Expression of collagen II in chondrocytes**.

(a) IHC staining of collagen II was performed on chondrocytes obtained from femur head cartilage from patients. Scale bar, 50 μm.

**Supplementary Figure 2. Intraperitoneal (IP) injection of curdlan induces inflammation in SKG mice**.

(a) Representative SKG mice 12 weeks after curdlan injection. Insets show a healthy, untreated SKG mouse (left) and a curdlan-treated SKG mouse with arthritis (right). (b) Clinical scores in curdlan-treated SKG mice and untreated SKG mice. Values are shown as the mean ± SD. **P < 0.01.

**Supplementary Figure 3. Cartilage structure in the hip joint from an AS patient was destroyed by granulation tissue**.

(a) H&E staining was performed on femur head cartilage from AS patients. Scale bar, 100 μm.

**Supplementary Figure 4. The ERS alleviator 4-PBA inhibited the angiogenic effect of chondrocytes**.

(a&b) Transwell migration assays were performed on HUVECs with CM (a). Migratory cells were calculated and quantified using ImageJ software (b). Scale bar, 100 μm. Values are shown as the mean ± SD. *P < 0.05, **P < 0.01.

**Supplementary Figure 5. ATF6 knockdown inhibited the angiogenic effect of chondrocytes**. (a&b) Transwell migration assays were performed on HUVECs with CM (a). Migratory cells were calculated and quantified using ImageJ software (b). Scale bar, 100 μm. Values are shown as the mean±S.D. *P<0.05, **P<0.01.

**Supplementary Figure 6. Knockdown of ATF6 expression in chondrocytes**.

(a&b) qRT-PCR and WB measurements of ATF6 expression in chondrocytes.

**Supplementary Figure 7. The ATF6 inhibitor Ceapin-A7 inhibited the angiogenic effect of chondrocytes**.

(a) The protein level of FGF2 in chondrocytes was measured after treatment with Ceapin-A7. (b) Transwell migration assays were performed on HUVECs with CM. The numbers of migratory cells were calculated and quantified using ImageJ software. Scale bar, 100 μm. Values are shown as the mean ± SD. *P < 0.05, **P < 0.01.

**Supplementary Figure 8. Ceapin-A7 inhibited ERS-induced FGF2 expression and osteogenesis in mice**.

(a) Diagram of the intra-articular injection of saline, Tg or Tg+Ceapin-A7 into C57BL/6 mice for 2 weeks. (b) IHC staining of ATF6, GRP78 and FGF2 was performed on knee joints from C57BL/6 mice after treatment with saline, Tg or Tg+Ceapin-A7 for 2 weeks. Scale bar, 50 μm. (c) IF staining of knee joints from C57BL/6 mice showing endothelial cells (CD31^+^) and activated osteoblasts (OCN^+^) around knee joints. Scale bar, 50 μm. Quantification of the proportions of CD31^+^ vessels and OCN^+^ cells.

**Supplementary Table S1.**
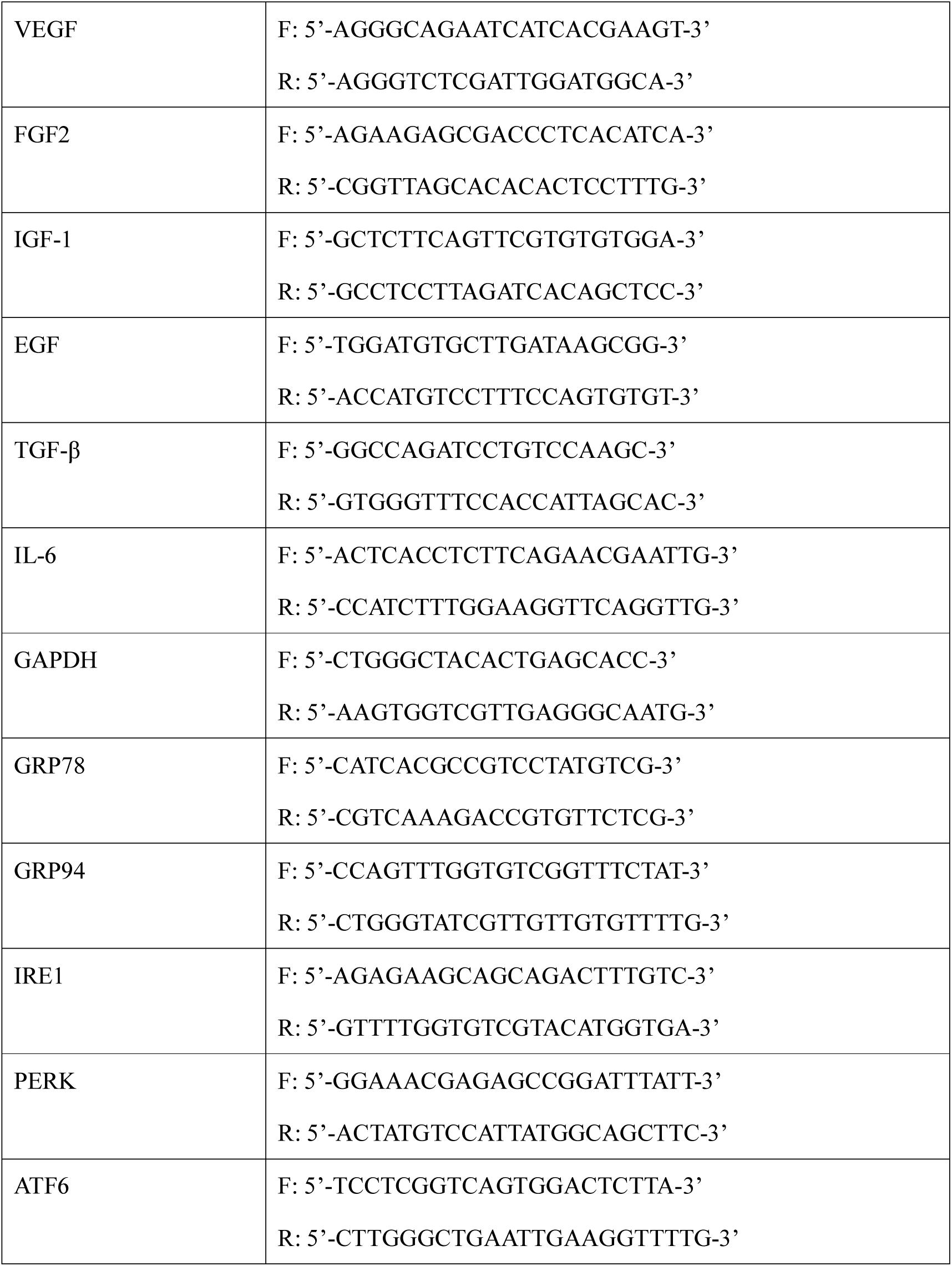
Primer pairs used for qRT-PCR.

## Notes

### Competing Interest Statement

The authors have declared no competing interest.

## References

1 Sieper J, Poddubnyy D. Axial spondyloarthritis. Lancet 2017; 390: 73–84.

2 Landewé R, Dougados M, Mielants H, van der Tempel H, van der Heijde D. Physical function in ankylosing spondylitis is independently determined by both disease activity and radiographic damage of the spine. Ann Rheum Dis 2009; 68: 863–7.

3 van der Heijde D, Landewé R, Baraliakos X, et al. Radiographic findings following two years of infliximab therapy in patients with ankylosing spondylitis. Arthritis Rheum 2008; 58: 3063–70.

4 van der Heijde D, Landewé R, Einstein S, et al. Radiographic progression of ankylosing spondylitis after up to two years of treatment with etanercept. Arthritis Rheum 2008; 58: 1324–31.

5 van der Heijde D, Salonen D, Weissman BN, et al. Assessment of radiographic progression in the spines of patients with ankylosing spondylitis treated with adalimumab for up to 2 years. Arthritis Res Ther 2009; 11: R127.

6 Poddubnyy D, Sieper J. Mechanism of new bone formation in axial spondyloarthritis. Curr Rheumatol Rep 2017; 19: 55.

7 Molnar C, Scherer A, Baraliakos X, et al. TNF blockers inhibit spinal radiographic progression in ankylosing spondylitis by reducing disease activity: results from the Swiss Clinical Quality Management cohort. Ann Rheum Dis 2018; 77: 63–9.

8 Baraliakos X, Haibel H, Listing J, Sieper J, Braun J. Continuous long-term anti-TNF therapy does not lead to an increase in the rate of new bone formation over 8 years in patients with ankylosing spondylitis. Ann Rheum Dis 2014; 73: 710–5.

9 Wendling D, Verhoeven F, Prati C. Anti-IL-17 monoclonal antibodies for the treatment of ankylosing spondylitis. Expert Opin Biol Ther 2019; 19: 55–64.

10 Lories RJ, Haroon N. Evolving concepts of new bone formation in axial spondyloarthritis: insights from animal models and human studies. Best Pract Res Clin Rheumatol 2017; 31: 877–86.

11 Lories RJ. Advances in understanding the pathophysiology of spondyloarthritis. Best Pract Res Clin Rheumatol 2018; 32: 331–41.

12 Krishnan Y, Grodzinsky AJ. Cartilage diseases. Matrix Biol 2018; 71-72: 51–69.

13 Bleil J, Sieper J, Maier R, et al. Cartilage in facet joints of patients with ankylosing spondylitis (AS) shows signs of cartilage degeneration rather than chondrocyte hypertrophy: implications for joint remodeling in AS. Arthritis Res Ther 2015; 17: 170.

14 Bleil J, Maier R, Hempfing A, Sieper J, Appel H, Syrbe U. Granulation tissue eroding the subchondral bone also promotes new bone formation in ankylosing spondylitis. Arthritis Rheumatol 2016; 68: 2456–65.

15 Bleil J, Maier R, Hempfing A, et al. Histomorphologic and histomorphometric characteristics of zygapophyseal joint remodeling in ankylosing spondylitis. Arthritis Rheumatol 2014; 66: 1745–54.

16 Binet F, Sapieha P. ER stress and angiogenesis. Cell Metab 2015; 22: 560–75.

17 Bettigole SE, Glimcher LH. Endoplasmic reticulum stress in immunity. Annu Rev Immunol 2015; 33: 107–38.

18 Hetz C, Papa FR. The unfolded protein response and cell fate control. Mol Cell 2018; 69: 169–81.

19 Ghosh R, Lipson KL, Sargent KE, et al. Transcriptional regulation of VEGF-A by the unfolded protein response pathway. PLoS One 2010; 5: e9575.

20 Hillary RF, FitzGerald U. A lifetime of stress: ATF6 in development and homeostasis. J Biomed Sci 2018; 25: 48.

21 Guo F, Han X, Wu Z, et al. ATF6a, a Runx2-activable transcription factor, is a new regulator of chondrocyte hypertrophy. J Cell Sci 2016; 129: 717–28.

22 Ranganathan V, Gracey E, Brown MA, Inman RD, Haroon N. Pathogenesis of ankylosing spondylitis – recent advances and future directions. Nat Rev Rheumatol 2017; 13: 359–67.

23 Sakaguchi N, Takahashi T, Hata H, et al. Altered thymic T-cell selection due to a mutation of the ZAP-70 gene causes autoimmune arthritis in mice. Nature 2003; 426: 454–60.

24 Ruutu M, Thomas G, Steck R, et al. β-glucan triggers spondylarthritis and Crohn’s disease-like ileitis in SKG mice. Arthritis Rheum 2012; 64: 2211–22.

25 Grootjans J, Kaser A, Kaufman RJ, Blumberg RS. The unfolded protein response in immunity and inflammation. Nat Rev Immunol 2016; 16: 469–84.

26 Gallagher CM, Walter P. Ceapins inhibit ATF6α signaling by selectively preventing transport of ATF6α to the Golgi apparatus during ER stress. Elife 2016; 5: e11880.

27 Kokame K, Kato H, Miyata T. Identification of ERSE-II, a new cis-acting element responsible for the ATF6-dependent mammalian unfolded protein response. J Biol Chem 2001; 276: 9199–205.

28 Cinaroglu A, Gao C, Imrie D, Sadler KC. Activating transcription factor 6 plays protective and pathological roles in steatosis due to endoplasmic reticulum stress in zebrafish. Hepatology 2011; 54: 495–508.

29 McInnes IB, Schett G. Cytokines in the pathogenesis of rheumatoid arthritis. Nat Rev Immunol 2007; 7: 429–42.

30 Haynes KR, Pettit AR, Duan R, et al. Excessive bone formation in a mouse model of ankylosing spondylitis is associated with decreases in Wnt pathway inhibitors. Arthritis Res Ther 2012; 14: R253.

31 Lories RJ, Derese I, Luyten FP. Modulation of bone morphogenetic protein signaling inhibits the onset and progression of ankylosing enthesitis. J Clin Invest 2005; 115: 1571–9.

32 Ruiz-Heiland G, Horn A, Zerr P, et al. Blockade of the hedgehog pathway inhibits osteophyte formation in arthritis. Ann Rheum Dis 2012; 71: 400–7.

33 Xie Z, Wang P, Li Y, et al. Imbalance between bone morphogenetic protein 2 and noggin induces abnormal osteogenic differentiation of mesenchymal stem cells in ankylosing spondylitis. Arthritis Rheumatol 2016; 68: 430–40.

34 Benjamin M, Toumi H, Suzuki D, Hayashi K, McGonagle D. Evidence for a distinctive pattern of bone formation in enthesophytes. Ann Rheum Dis 2009; 68: 1003–10.

35 Pedersen SJ, Maksymowych WP. The pathogenesis of ankylosing spondylitis: an update. Curr Rheumatol Rep 2019; 21: 58.

36 Dong W, Zhang Y, Yan M, Liu H, Chen Z, Zhu P. Upregulation of 78-kDa glucose-regulated protein in macrophages in peripheral joints of active ankylosing spondylitis. Scand J Rheumatol 2008; 37: 427–34.

37 Ciccia F, Accardo-Palumbo A, Rizzo A, et al. Evidence that autophagy, but not the unfolded protein response, regulates the expression of IL-23 in the gut of patients with ankylosing spondylitis and subclinical gut inflammation. Ann Rheum Dis 2014; 73: 1566–74.

38 De Palma M, Biziato D, Petrova TV. Microenvironmental regulation of tumour angiogenesis. Nat Rev Cancer 2017; 17: 457–74.

39 Woodbury ME, Ikezu T. Fibroblast growth factor-2 signaling in neurogenesis and neurodegeneration. J Neuroimmune Pharmacol 2014; 9: 92–101.

40 Akl MR, Nagpal P, Ayoub NM, et al. Molecular and clinical significance of fibroblast growth factor 2 (FGF2 /bFGF) in malignancies of solid and hematological cancers for personalized therapies. Oncotarget 2016; 7: 44735–62.

41 Bossé Y, Rola-Pleszczynski M. FGF2 in asthmatic airway-smooth-muscle-cell hyperplasia. Trends Mol Med 2008; 14: 3–11.

42 Poudel SB, Min CK, Lee JH, et al. Local supplementation with plant-derived recombinant human FGF2 protein enhances bone formation in critical-sized calvarial defects. J Bone Miner Metab 2019; 37: 900–12.

43 Huang G, Zhao G, Xia J, et al. FGF2 and FAM201A affect the development of osteonecrosis of the femoral head after femoral neck fracture. Gene 2018; 652: 39–47.

44 Yang W, Cao Y, Zhang Z, et al. Targeted delivery of FGF2 to subchondral bone enhanced the repair of articular cartilage defect. Acta Biomater 2018; 69: 170–82.

45 Wang J, Liu S, Li J, Yi Z. The role of the fibroblast growth factor family in bone-related diseases. Chem Biol Drug Des 2019; 94: 1740–9.

46 Yu X, Qi Y, Zhao T, et al. NGF increases FGF2 expression and promotes endothelial cell migration and tube formation through PI3K/Akt and ERK/MAPK pathways in human chondrocytes. Osteoarthritis Cartilage 2019; 27: 526–34.

47 Kawane T, Qin X, Jiang Q, et al. Runx2 is required for the proliferation of osteoblast progenitors and induces proliferation by regulating Fgfr2 and Fgfr3. Sci Rep 2018; 8: 13551.

48 Lee MN, Kim JW, Oh SH, Jeong BC, Hwang YC, Koh JT. FGF2 stimulates COUP-TFII expression via the MEK1/2 pathway to inhibit osteoblast differentiation in C3H10T1/2 cells. PLoS One 2016; 11: e0159234.

