## Supplementary figures and images for "ATF6 aggravates angiogenesis-osteogenesis coupling during ankylosing spondylitis by mediating FGF2 expression in chondrocytes"

### Supplemental Figure 1

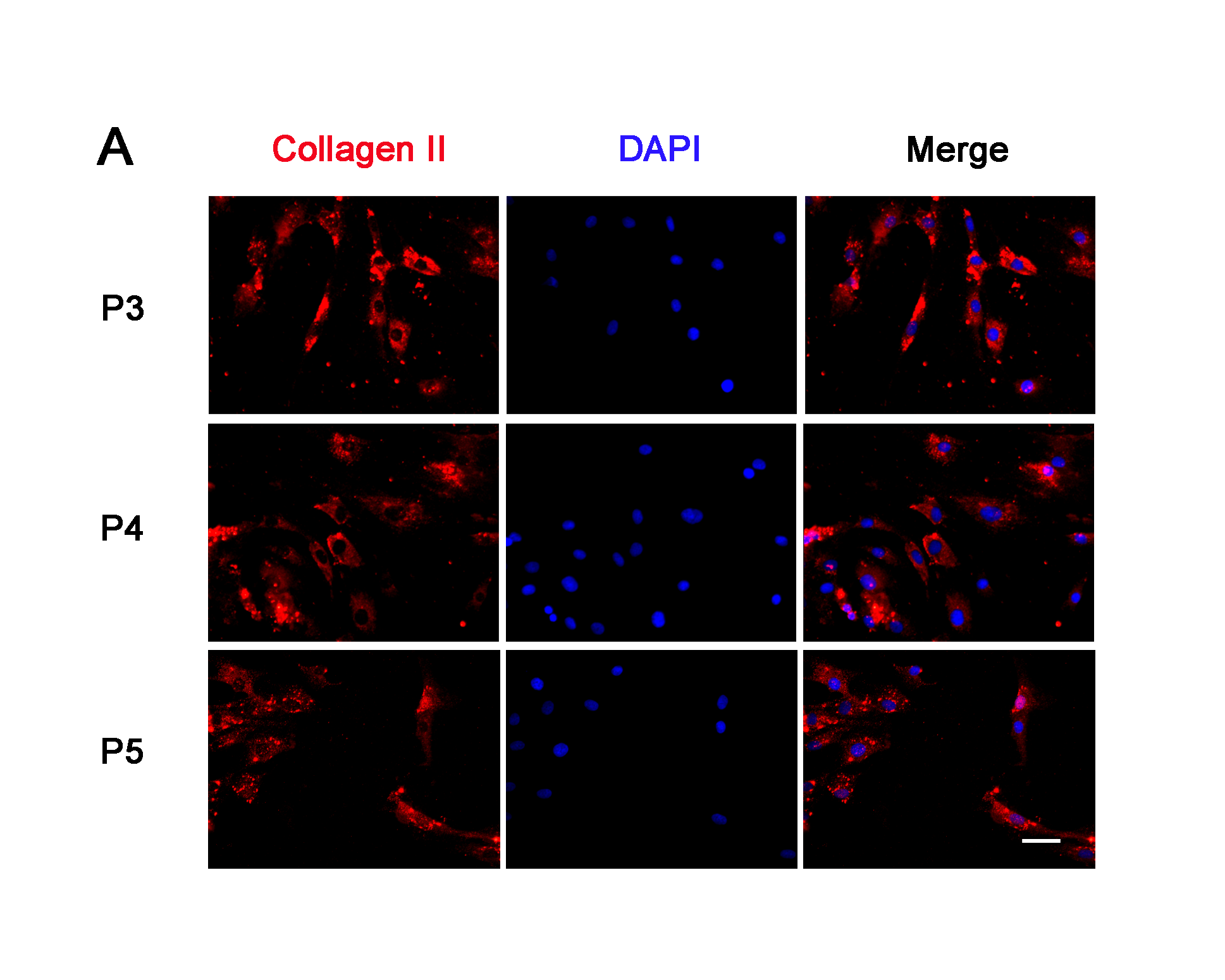

### Supplemental Figure 2

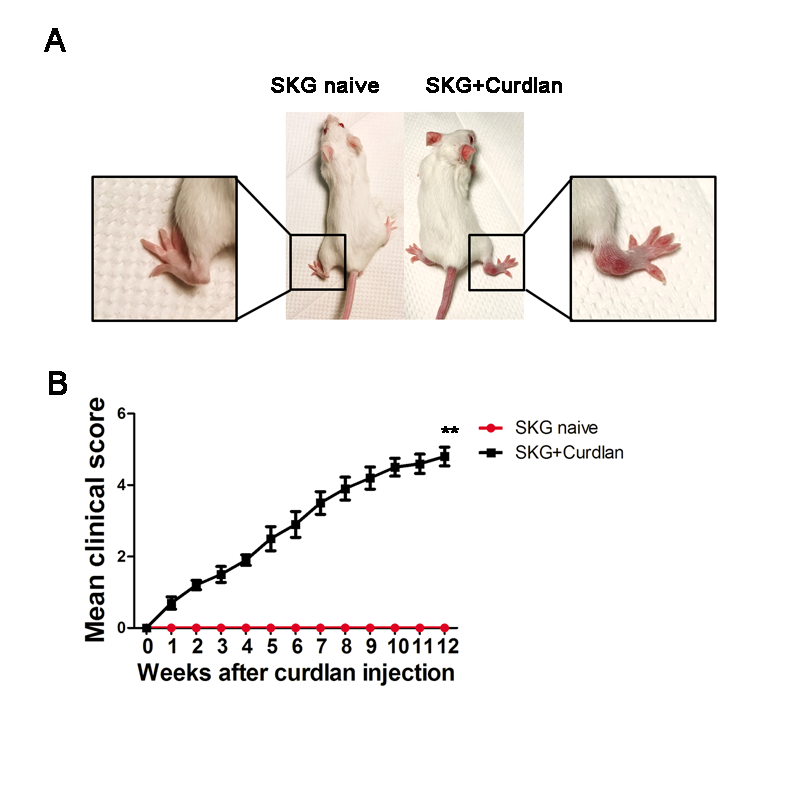

### Supplemental Figure 3

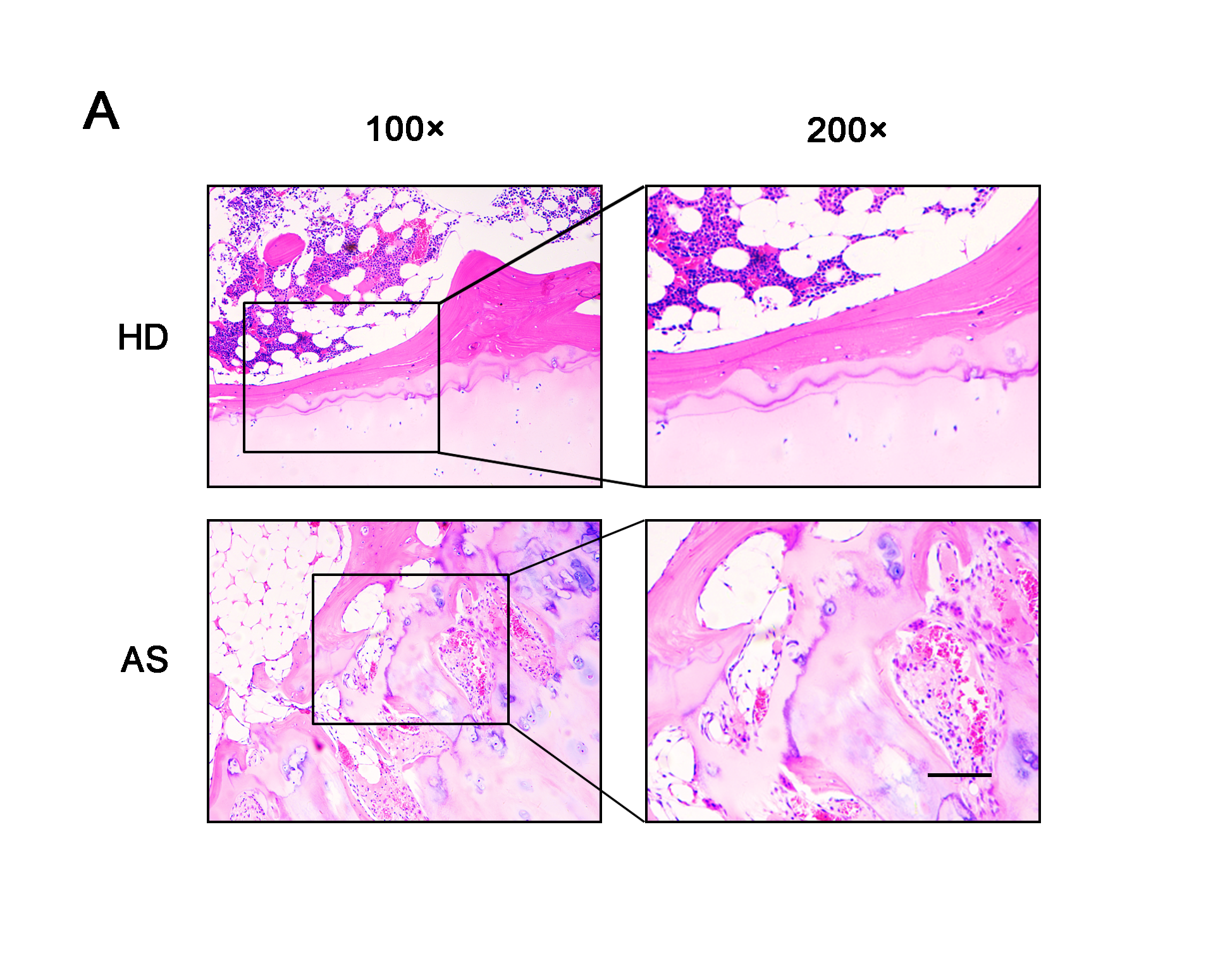

### Supplemental Figure 4

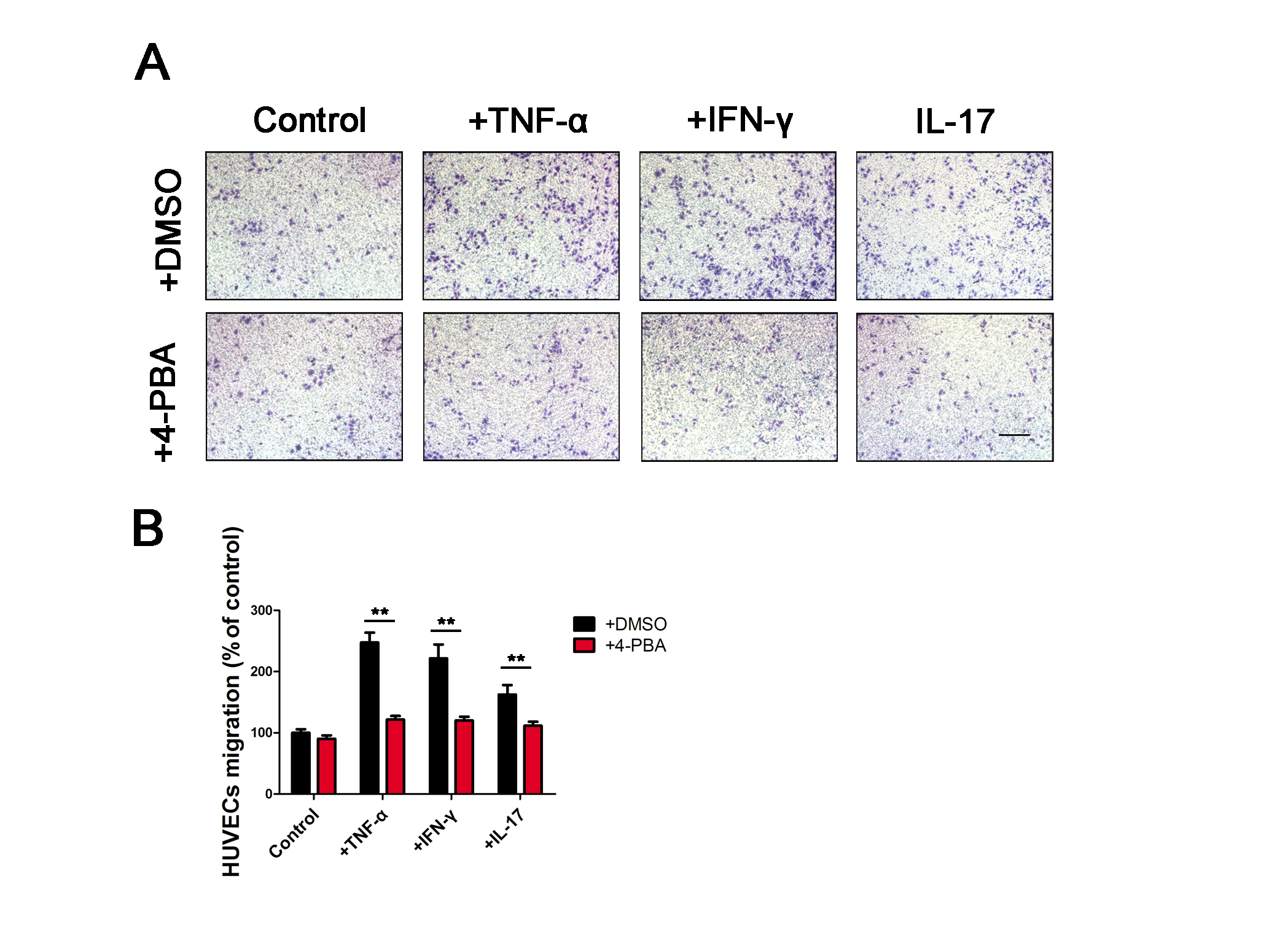

### Supplemental Figure 5

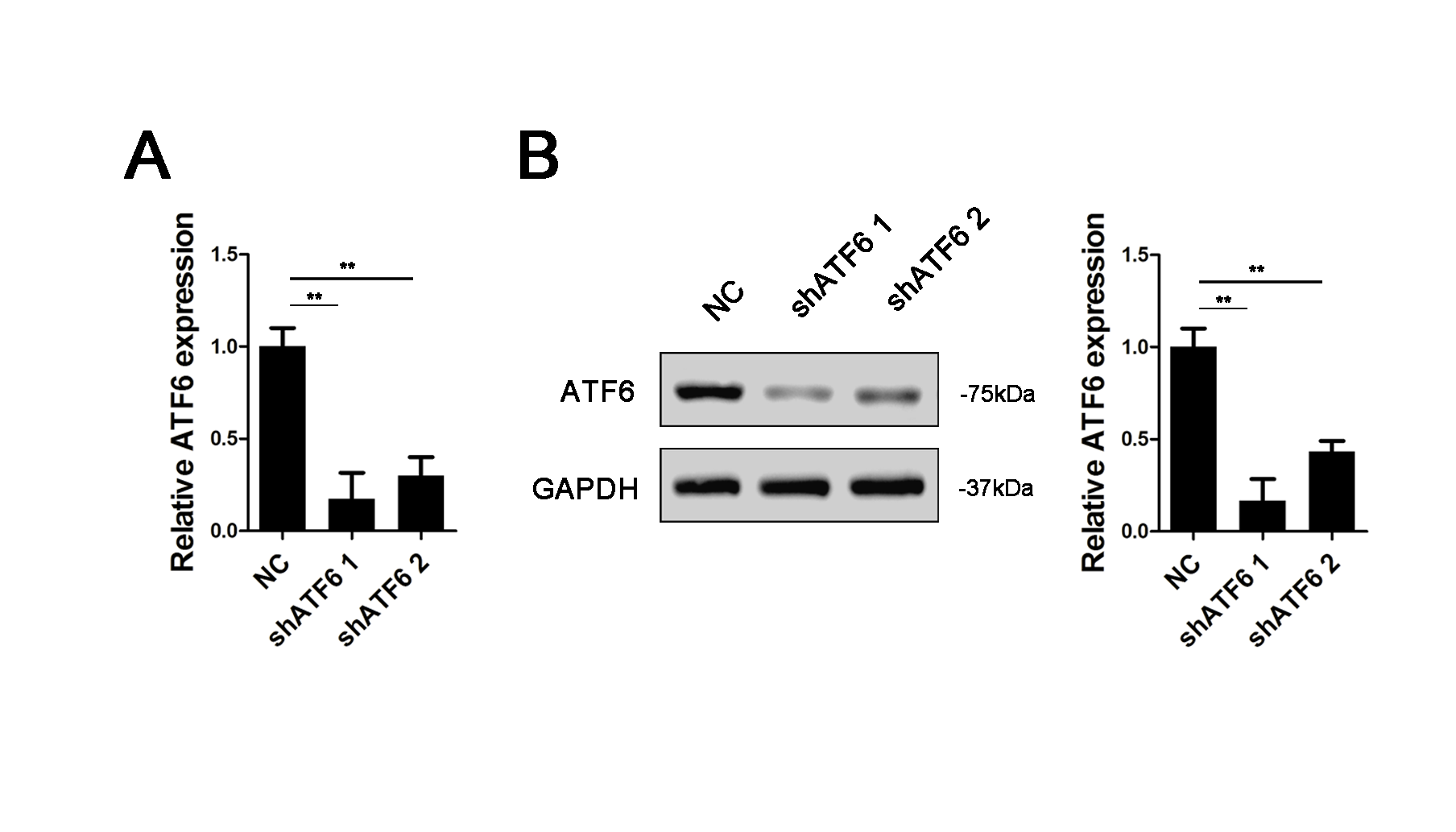

### Supplemental Figure 6

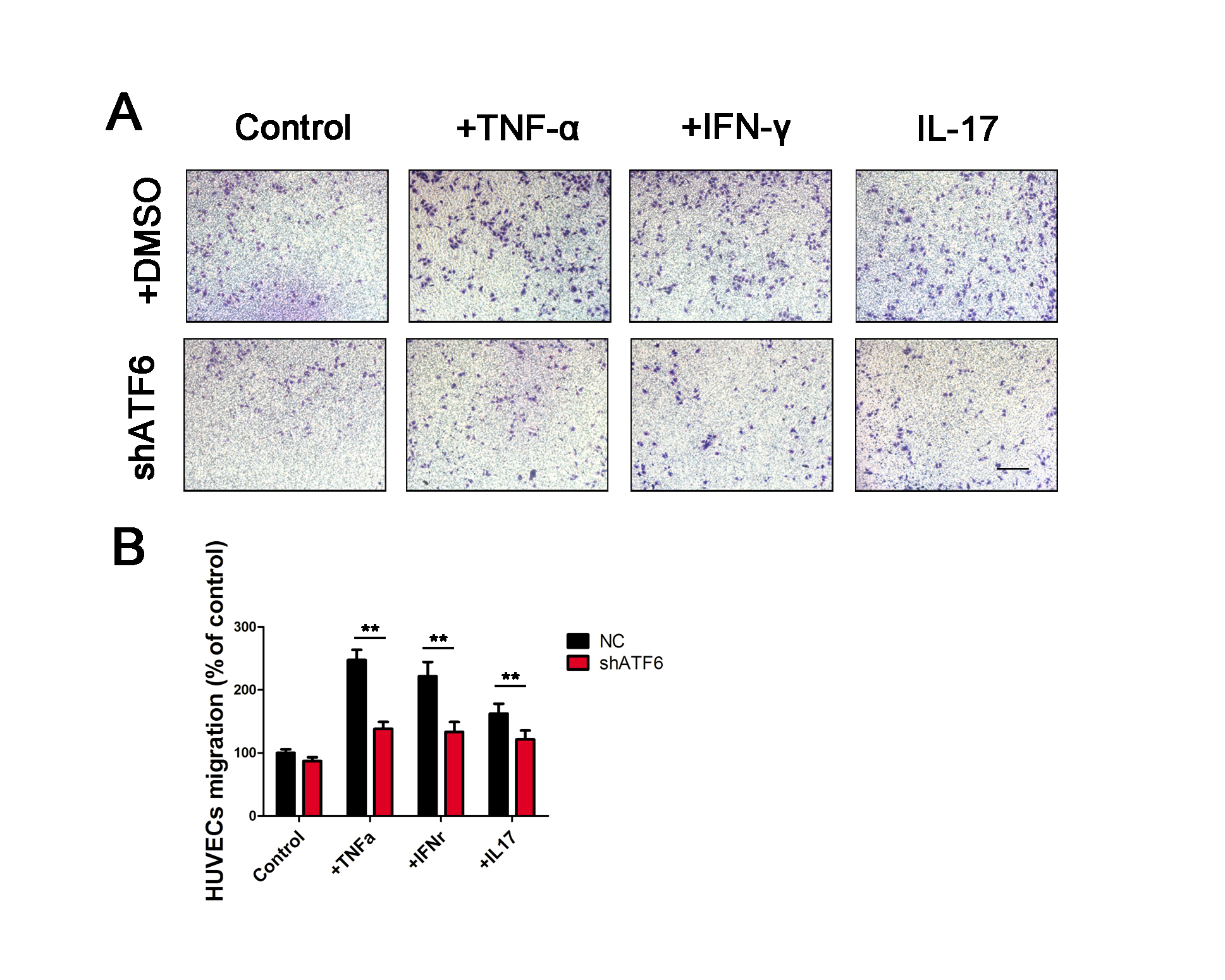

### Supplemental Figure 7

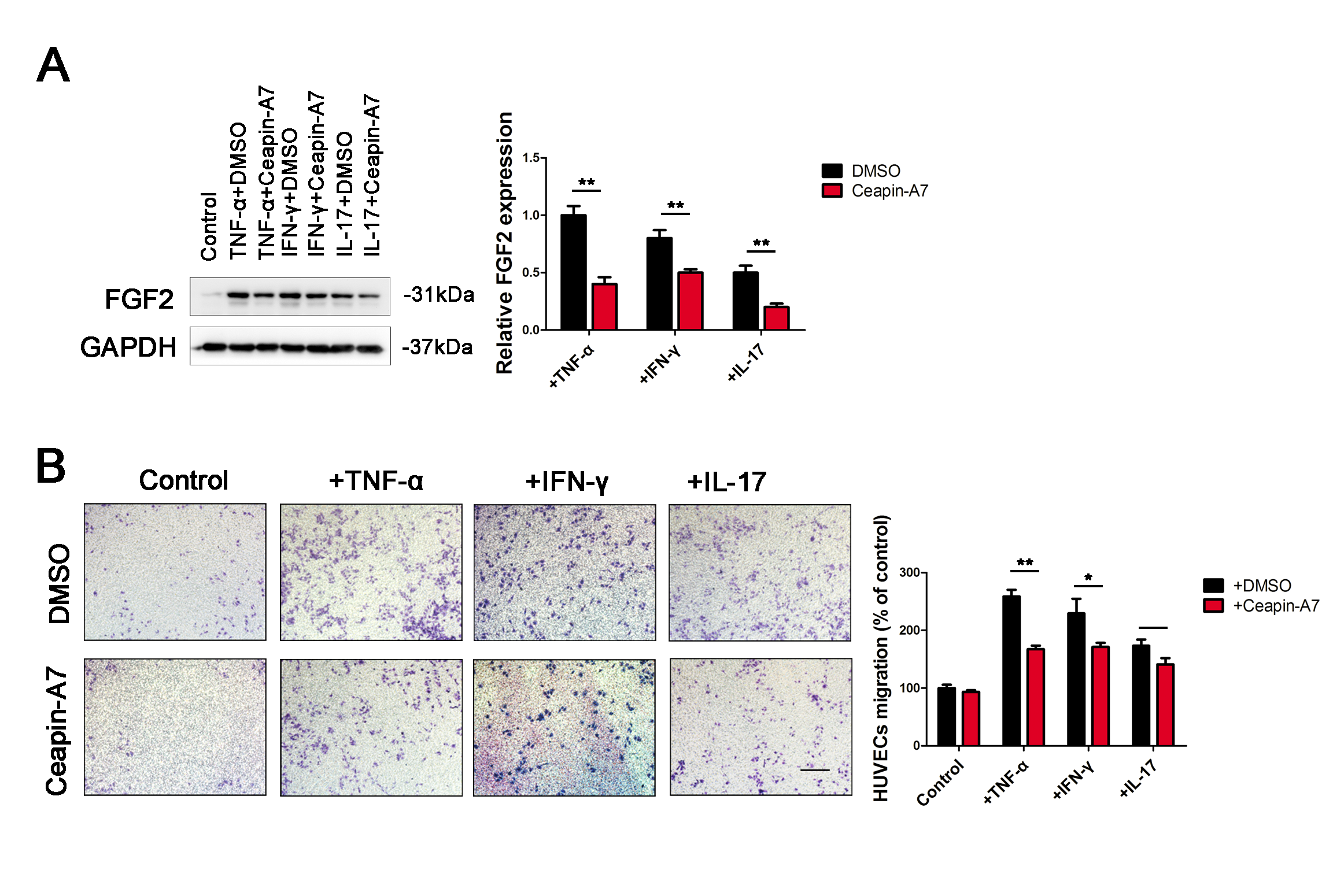

### Supplemental Figure 8

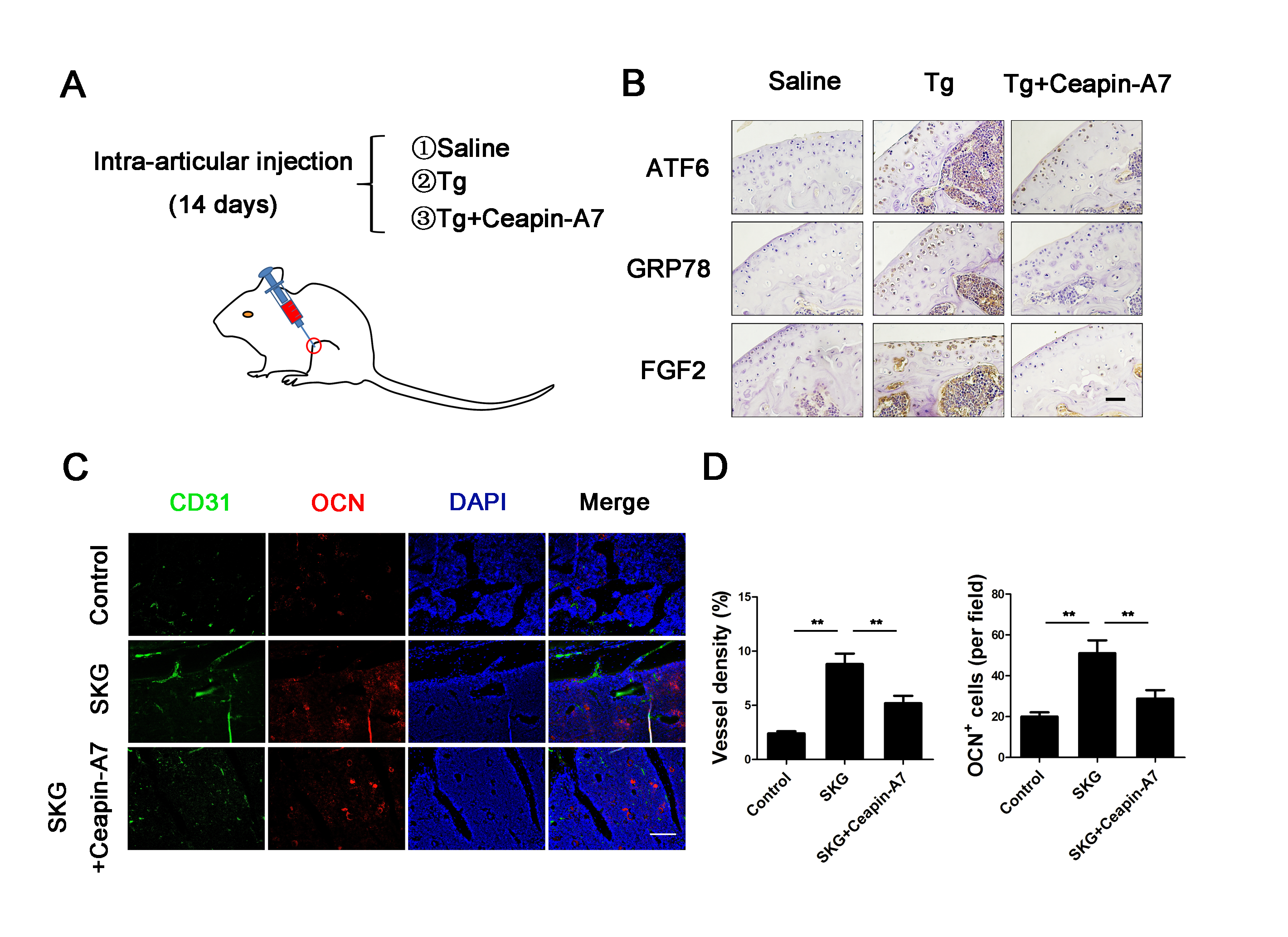
